# Epigenetic insights into GABAergic development in Dravet Syndrome iPSC and therapeutic implications

**DOI:** 10.1101/2023.10.10.561691

**Authors:** Jens Schuster, Xi Lu, Yonglong Dang, Joakim Klar, Amelie Wenz, Niklas Dahl, Xingqi Chen

## Abstract

Dravet syndrome (DS) is a devastating early onset refractory epilepsy syndrome caused by variants in the *SCN1A* gene. A disturbed GABAergic interneuron function is implicated in the progression to DS but the underlying developmental and pathophysiological mechanisms remain elusive, in particularly at the chromatin level. In this study, we utilized induced pluripotent stem cells (iPSCs) derived from DS cases and healthy donors to model disease-associated epigenetic abnormalities of GABAergic development. Employing the ATAC-Seq technique, we assessed chromatin accessibility at multiple time points (Day 0, Day 19, Day 35, and Day 65) of GABAergic differentiation. Additionally, we elucidated the effects of the commonly used anti-seizure drug valproic acid (VPA) on chromatin accessibility in GABAergic cells. The distinct dynamics in chromatin profile of DS iPSC predicted accelerated early GABAergic development, evident at D19, and diverged further from the pattern in control iPSC with continued differentiation, indicating a disrupted GABAergic maturation. Exposure to VPA at D65 reshaped the chromatin landscape at a variable extent in different iPSC-lines and rescued the observed dysfunctional development in some DS iPSC-GABA. This study provides the first comprehensive investigation on the chromatin landscape of GABAergic differentiation in DS-patient iPSC, offering valuable insights into the epigenetic dysregulations associated with interneuronal dysfunction in DS. Moreover, our detailed analysis of the chromatin changes induced by VPA in iPSC-GABA holds the potential to improve development of personalized and targeted anti-epileptic therapies.

## Introduction/Background

Dravet syndrome (DS) is an early onset and intractable epilepsy with an unfavorable long-term outcome. The first seizures are usually triggered by fever within the first 12 months of life^1,2^. Characteristic clinical features are age-related progression of seizures, cognitive decline, behavioral problems and movement disorder. The complex neurological symptoms associated with DS suggest underlying pathophysiological mechanisms that interfere with brain development^1–4^. Approximately 80% of DS cases carry heterozygous variants in the *SCN1A* gene encoding the α-subunit voltage-gated sodium channel (Nav1.1)^1,5^. Prior insights on the neuropathophysiology in DS have come from the *Scn1a* heterozygous mice, indicating a vital role of cortical interneurons in the pathogenesis. Mice with Nav1.1 haploinsufficiency in GABAergic interneurons exhibit spontaneous seizures and behavioral abnormalities^6–9^ that are associated with subnormal sodium currents and impaired excitability of inhibitory interneurons^10,11^. Additionally, human iPSC derived neural cells carrying distinct pathogenic *SCN1A* variants have recapitulated electrophysiological and molecular perturbations in DS^12–16^. In combination, these studies have confirmed that *SCN1A* variants cause delayed sodium currents in activated and mainly inhibitory GABAergic interneurons, supporting the notion that seizures in DS are caused by deficient cortical inhibition^12^.

Interneuronal progenitors emerge from the embryonic subpallium, mainly the median ganglionic eminence, and migrate tangentially to cortical progenitor zones. The second half of gestation is a period of rapid development of the human cortical GABAergic system that continues after term^17,18^. The extended developmental processes into mature GABAergic interneurons require continuous and complex remodeling of the chromatin structure for transcriptional adaptations^19^. While it is now clear that the distinct features in DS is associated with a reduced sodium current density in Nav1.1 haploinsufficient interneurons and a disinhibition of the cortical network^6,13,20^, the underlying epigenetic mechanisms are poorly understood.

Treatment of DS patients is challenging and seizure control is rarely attainable^21^. The most frequently used first-line drug is valproic acid (VPA)^1^. However, more than 50% of DS patients are drug resistant without any decrease in seizure-frequency upon treatment^1^, indicating endogenous and subject-specific responses to VPA. The mechanisms by which VPA act are not fully understood, but studies suggest that the drug inhibits histone deacetylase (HDAC) with an impact on chromatin structure and gene expression^22,23^. Furthermore, a direct interaction of VPA with sodium- or calcium-gated ion channels (e.g. Nav1.1) and enzymes crucial for GABA turnover has been proposed^22^. While these studies have brought important information on the mode of action of VPA^24–28^, the specific effects of VPA on chromatin architecture of inhibitory GABAergic interneurons have not yet been investigated to our knowledge.

We recently established induced pluripotent stem cell (iPSC) lines carrying distinct disease-causing variants in the *SNC1A* gene and modeled GABAergic interneuron development in DS^13,29^. The DS-patient iPSC derived GABAergic neurons recapitulated electrophysiological abnormalities and gene expression analysis showed enrichment for histone modifications, suggesting epigenetic abnormalities^13^. We therefore sought to explore the dynamics of open chromatin in our DS-model of GABAergic development in DS using the assay of transposase accessible chromatin sequencing (ATAC-Seq). The methodology has previously been used to study chromatin accessibility of annotated genes during brain development^30–32^ and, specifically, in interneuronal differentiation^19,33^. Herein, we employed ATAC-Seq to investigate the dynamics of chromatin accessibility in iPSC of DS-patients and healthy donors at four different time points (Day 0, Day 19, Day 35, and Day 65) of GABAergic development. Moreover, we explored the effect of VPA on chromatin accessibility in GABAergic neurons. Our study uncovered accelerated chromatin changes in DS patient cells during the initial phase of GABAergic development (up to Day 19) when compared to cells of healthy donors. Further differentiation revealed that DS-patient GABAergic cells acquire a distinct chromatin profile when compared to that of control cells. Notably, VPA treatment of GABAergic neurons leads to unspecific genome-wide changes in chromatin architecture that predicts a promoted GABAergic development in some iPSC lines. This study represents the first investigation of the chromatin dynamics of GABAergic development in DS-patient iPSC, shedding light on the underlying epigenetic abnormalities. Moreover, the in-depth characterization of chromatin changes in GABAergic neurons induced by VPA may bring important information for the development of individualized anti-seizure therapies.

## Results

### Dynamic chromatin accessibility in an iPSC model of GABAergic development

We previously established a protocol for GABAergic interneuronal differentiation of iPSC^13,29^. Analysis of RNAseq data at D19 and D65 of GABAergic differentiation revealed *SCN1A* expression in neuronal cells derived from healthy donors and from patients with Dravet disease (**Supplementary Figure 1A**). The *SCN1A* expression was confirmed with qPCR at D19, D35, and D65 of differentiation whereas no expression could be detected in undifferentiated iPSC (D0; **Supplementary Figure 1B**). We then applied ATAC-Seq to investigate the dynamics of chromatin accessibility in GABAergic development associated with *SCN1A* variants. We first obtained a reference for chromatin accessibility changes during GABAergic interneuronal differentiation in iPSC from three healthy donors (Ctl) using ATAC-Seq^35^ (**Figure 1A**). The differentiating cultures were collected at four distinct time points of differentiation at day 0 (D0; termed iPSC), D19 (neural progenitor cells; NPC), D35 (intermediate neuronal cells; imN) and D65 (GABAergic interneurons; GABA). The cultures displayed expression of marker genes confirming the development into GABAergic interneurons^13^. As previously shown, cultures stained positive for pluripotent stem cell markers such as NANOG, SOX2, and OCT4 at D0. The expression of these markers decreased upon neural induction and disappeared with further differentiation. Markers for neural progenitor cells (FOXG1, PAX6) were expressed at D19 followed by the expression of immature neuronal cell markers (DCX and NKX2.1) at D35 and of markers for GABAergic interneurons (GAD1, TUBB3) at D65 (**Supplementary Figure 1C and Schuster et al. 2019**^13^).

**Figure 1:**
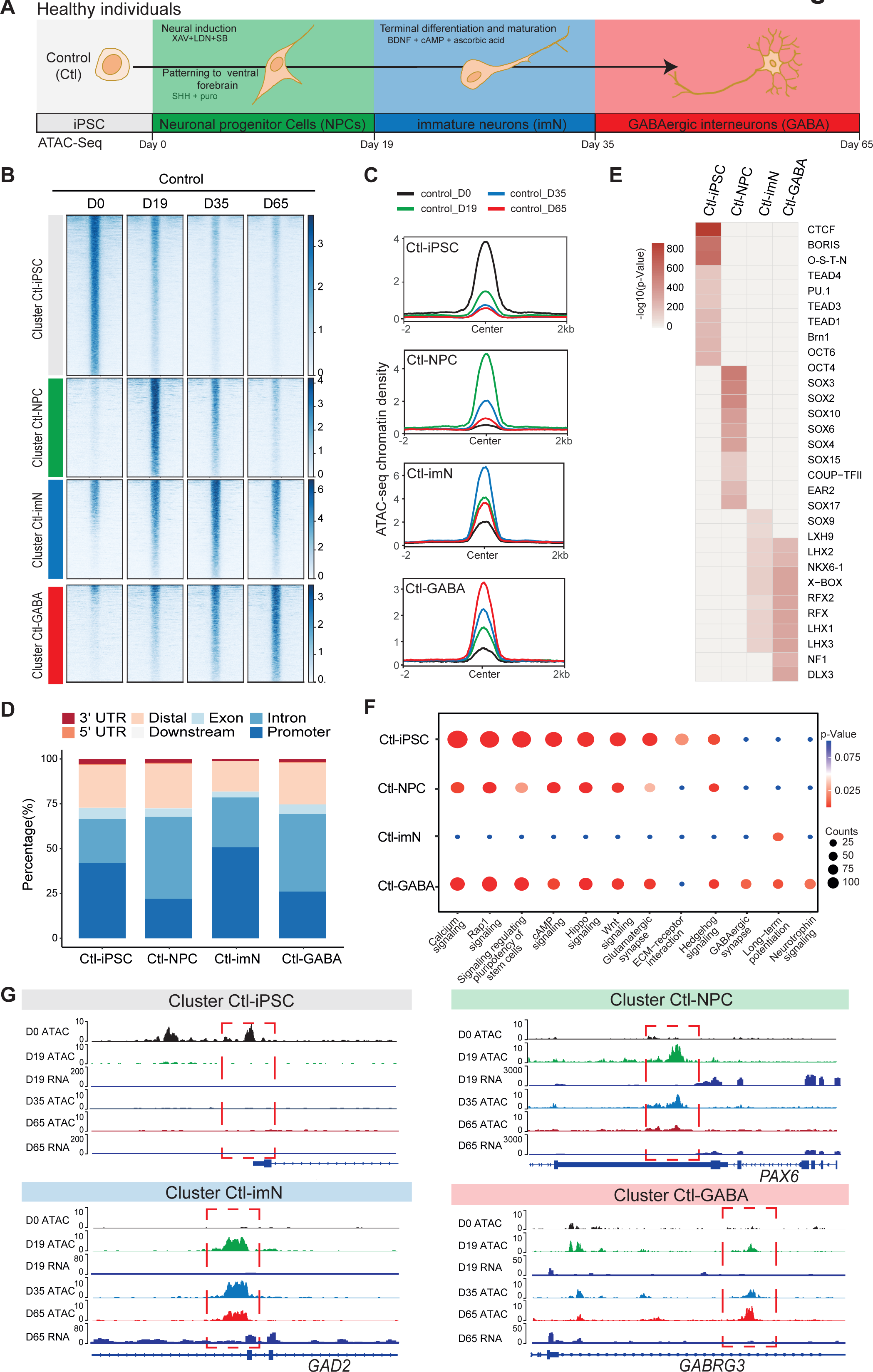
Chromatin accessibility dynamics during GABAeric interneuron differentiation of control iPSCs. **A.** Schematic illustration of GABAergic interneuron differentiation protocol. Cells from iPSC differentiation were collected for ATAC-Seq. Interneuron differentiation was grouped into four stages: iPSC (Day 0), neural progenitor cells; NPCs (Day 19), immature neurons, imN (Day 35), GABAergic interneurons, GABA (Day 65). **B.** Heatmap of four Clusters identified by time series analysis using temporal changes in chromatin accessibility compared to D0 (Day 0) and other points. The signal strengths were denoted by color intensities. **C.** Line plots showing chromatin accessibility at cluster-specific regions from each time point. **D.** Barplot for genomic distribution of differential chromatin accessible regions for each cluster. **E.** Heatmap for top 10 TFs enrichment in each cluster. The significance was denoted by color intensities. O-S-E-N = OCT4-SOX2-TCF-NANOG. **F.** Bubble plot of KEGG pathway enrichment for each cluster. p-value and enrichment were indicated. The corresponding comprehensive list of enrichment terms can be found in Supplementary Table 2. **G.** Genome browser view showing representative differential chromatin accessible regions at the indicated gene loci. Additionally, RNA-seq data^13^ is visualized at D19 and D65.

The quality of ATAC-Seq data from each sample was validated by sequencing depth comparison, genome-wide correlation between technical replicates and by calculating the fraction of reads in enriched peaks (FRiP), the transcription start site enrichment score (**Supplementary Figure 2A**-D). We captured accessible chromatin peaks at each time-point of differentiation: 58,382 peaks at D0 (Ctl-iPSC), 68,002 peaks at D19 (Ctl-NPC), 52,933 peaks at D35 (Ctl-iMN), and 74,561 peaks at D65 (Ctl-GABA) (**Supplementary Table 1**). Genomic annotation of the accessible chromatin sites with respect to promoters, introns and exons revealed a heterogenous distribution in iPSC, NPC, imN and GABA (**Supplementary Figure 2E).** Furthermore, principal component analysis (PCA) of chromatin accessibility at the four time points demonstrated that the three biological replicates separated into four clusters corresponding to each of the four time points of GABAergic development (**Supplementary Figure 2F**), albeit with some degree of variability at the intermediate time points (D19 (NPC) and D35 (imN)), possibly reflecting cell line specific and endogenous differences reported previously^36^. We therefore treated the three Ctl-iPSC-lines as biological replicates for downstream analysis. We also performed genome-wide correlation of ATAC-Seq with our previously published RNA-Seq data at D19 and D65 (**Supplementary Figure 2G**). The correlation ranged between 0.52 and 0.57, further indicating the good quality of the ATAC-Seq data.

We then characterized changes in chromatin accessibility during GABAergic development in Ctl-iPSC. The ATAC-Seq peaks in iPSC (D0) were used as reference and compared to the accessible peaks in NPC, imN and GABA. To avoid capturing the dynamic changes of accessible regions caused by variability across individuals, we initially compared the dynamic changes of chromatin accessibility cell line by cell line across differentiation. Subsequently, we extracted the common changes observed across different cell lines at each time point (**Methods**). In total, we identified 19,931 significant differential peaks (|log2(fold change (FC))|> 1, false discovery rate (FDR) < 0.01) (**Supplementary Figure 3A, Methods**). Next, we employed an unbiased cluster method to investigate the dynamics of these differential accessible peaks, resulting in the identification of four distinct clusters (Cluster 1-4) (**Figure 1B, Supplementary Figure -3B-D**). Cluster 1 (n=9,565) consisted of peaks that were open D0 (cluster Ctl-iPSC) and that gradually closed during differentiation. Cluster 2 (n=3,285) contained peaks that were predominantly open at D19 (cluster Ctl-NPC). Cluster 3 (n=3,533) exhibited a high chromatin accessibility at both D19 and D35 (cluster Ctl-imN) but more closed at D65. Cluster 4 (n=3,545) displayed the highest accessibility specifically at D65 (cluster Ctl-GABA; **Figure 1B, 1C**). A genomic annotation of the differential peaks in the four clusters with respect to promoters, exons, introns and distal regulatory sequences revealed differences mainly in introns and promoter regions. The Ctl-NPC cluster and Ctl-GABA cluster showed the highest proportion of open chromatin in introns (45.6% and 43.4%, respectively) whereas in the Ctl-iPSC and Ctl- imN clusters most accessible peaks were located in promoter regions (41.9% and 50.8%, respectively; **Figure 1D**).

We then performed transcription factor (TF) enrichment analysis on the differential peaks specific to each cluster (**Figure 1E**). The Ctl-iPSC cluster showed specific enrichment for motifs of TFs for pluripotency, such as BRN1, OCT6, among others (**Figure 1E**), whereas motifs for the SOX family of TFs were enriched in the Ctl-NPC-cluster. Ctl-NPC-specific transcription factor motifs (e.g., SOX family TFs) were detected in the NPC-specific cluster. Notably, motifs for common TFs (e.g., LHX family TFs, NKX6, RFX) as well as unique TFs

(e.g. SOX9, NF1 and DLX3) were enriched in both the Ctl-imN-specific cluster and the Ctl- GABA specific cluster. The SOX family of TFs are important regulators of neuronal development^37^ and DLX factors function in orchestrating transcriptional activity during cortical GABAergic development^38^. The temporal pattern of accessible and enriched TF motifs in our differentiating Ctl-iPSC cultures is thus consistent with the TFs required for neuronal and cortical GABAergic development *in vivo*, confirming our model system to be relevant.

Furthermore, we examined the biological processes associated with each cluster by assigning each differential peak to the nearest gene. The annotated genes were then used for KEGG analysis of each cluster (**Figure 1F, Supplementary Table 2**). In the Ctl-iPSC cluster we identified enrichment of pathways related to pluripotency, such as signaling regulating pluripotency of stem cells, while the Ctl-GABA cluster showed exclusive enrichment for pathways for GABAergic neurons, such as GABAergic synapse and Neurotrophin signaling (**Figure 1F**). Genome traces of annotated genes in each cluster validated the differential accessible chromatin, e.g. for *NANOG* in the Ctl-iPSC cluster, *PAX6* and *GAD2* in the Ctl-NPC and Ctl-imN clusters, respectively, and *GABRG3* in the Ctl-GABA cluster (**Figure 1G**). Gene expression analysis of RNA-seq data at D19 and D65, as well as with qPCR at D0, D19, D35, and D65, indicated that expression levels of the four genes was consistent with the chromatin accessibility patterns (**Supplementary Figure 3E**). Taken together, these findings show that the captured chromatin dynamics comply with the induction and development of GABAergic neurons.

### Dynamic chromatin accessibility in a DS-iPSC model of GABAergic development

Next, to gain a deeper understanding of chromatin dynamics during GABAergic development associated with DS, we used the same strategy as above for differentiation and ATAC-Seq analysis of iPSC-lines derived from the three DS patients (**Figure 2A, Supplementary** Figure 4). Unfortunately, one of the DS-iPSC lines (DD4A) did not differentiate up to D65 for unknown reasons why data at this specific time-point are based on two DS-iPSC lines.

**Figure 2:**
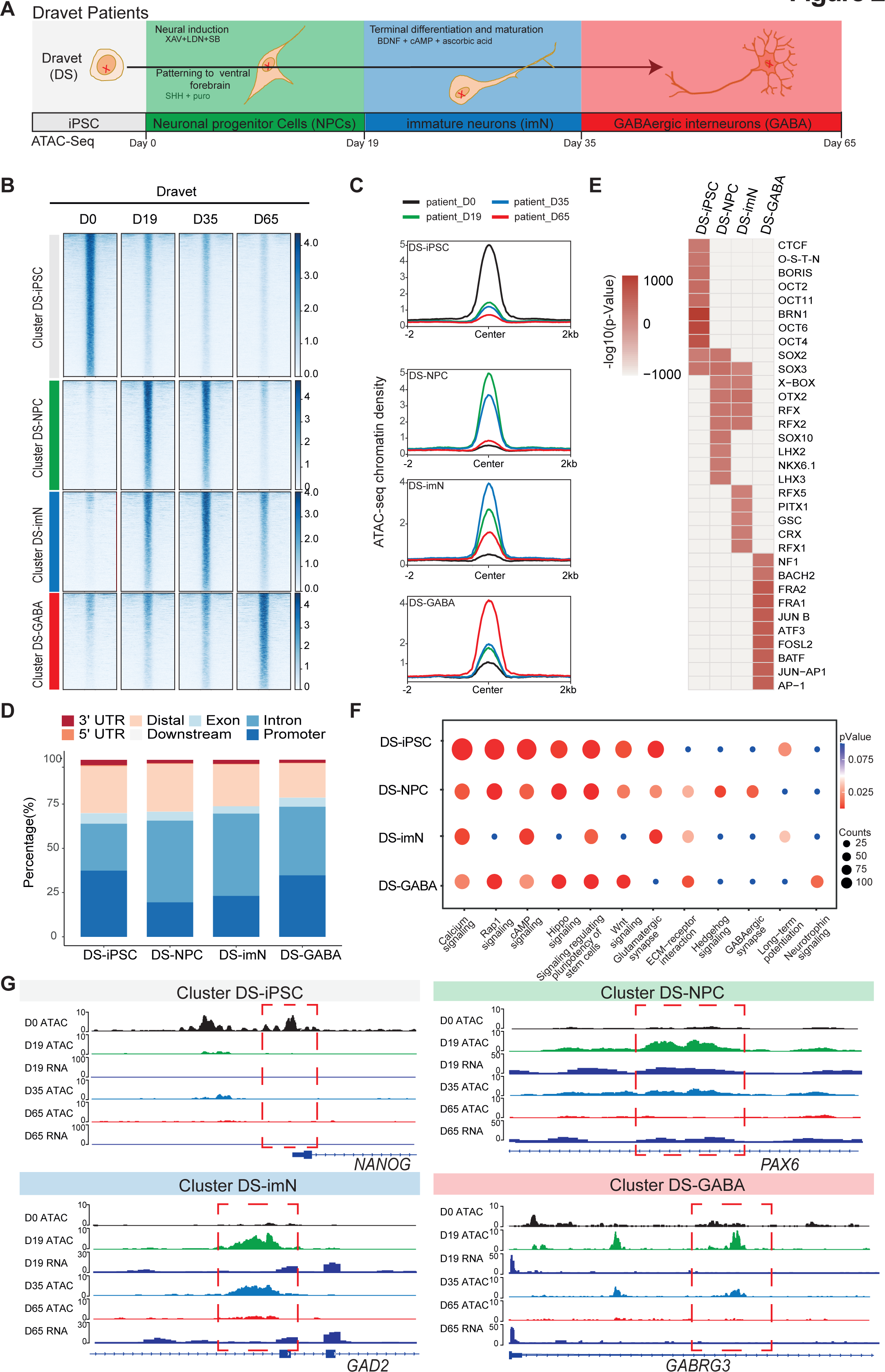
Chromatin accessibility dynamics during GABAergic differentiation of Dravet Syndrome patient iPSCs **A.** Schematic Illustration of the design for GABAergic differentiation in Dravet syndrome patients. Cells from iPSC differentiation were collected for ATAC-Seq. GABAergic differentiation in Dravet Syndrome patients was grouped into four stages: iPSC (Day 0), NPCs (Day 19), imN (Day 35), GABA(Day 65). **B.** Heatmap of four clusters identified by time series analysis using temporal changes in chromatin accessibility compared D0 (Day 0) and other points. The signal strengths were denoted by color intensities **C.** Line plots showing chromatin accessibility at cluster-specific regions from each time point. **D.** Barplot for genomic distribution of differential chromatin accessible regions for each cluster. **E.** Heatmap for top 10 TFs enrichment in each cluster. The significance was denoted by color intensities. O-S-E-N = OCT4-SOX2-TCF-NANOG. **F.** Bubble plot of KEGG pathway enrichment for each cluster. p-value and enrichment were indicated. The corresponding comprehensive list of enrichment terms can be found in Supplementary Table 4. **G.** Genome browser view showing representative differential chromatin accessible regions at the indicated gene loci. Additionally, RNA-seq data^13^ is visualized at D19 and D65.

The ATAC-Seq data showed high quality and reproducibility at all four time-points (**Supplementary Figure 4A**-D). In total, we detected 89,363 peaks in DS-iPSC, 79,270 peaks in DS-NPC, 113,519 peaks in DS-imN, and 41,665 peaks in DS-GABA (**Supplementary Table 3**). The genomic distribution of these accessible chromatin peaks with respect to promoters, introns and exons showed deviations when compared to Ctl-iPSC lines, mainly at D35 (DS-imN) and at D65 (DS-GABA) (**Supplementary Figure 4E**). Furthermore, the PCA of chromatin accessibility in DS-iPSC replicates showed that they clustered together at D0, D19 and D35 (**Supplementary Figure 4F**). Notably, the clusters at D19 (iPSC-NPC) and at D35 (iPSC-imN) showed a strong overlap in contrast to the well distributed clusters at all four time points in Ctl-iPSC lines (**Supplementary Figure 2F** and 4F). Genome-wide correlation of ATAC-Seq and RNA-seq data at D19 and D65 showed good correlation (0.49 to 0.55; **Supplementary Figure 4G**). The different dynamics of chromatin accessibility of DS-iPSC lines when compared to that of Ctl-iPSC lines thus suggest a disrupted developmental trajectory into DS-GABAergic interneurons consistent with an altered function of inhibitory GABAergic interneurons in *Scn1a* heterozygous mice.

To further characterize chromatin changes in GABAergic differentiation of DS-iPSC lines, we used the same strategy as for Ctl-iPSC lines by using the profile of chromatin accessibility in DS-iPSCs (D0) as a reference. To avoid capturing the dynamic changes of accessible regions caused by variability across individuals, we applied the same strategy as described for the control samples (Methods). In total, we identified 19,896 differential peaks (|log2(FC)| > 1, FDR < 0.01) (**Methods**) that were clustered into four groups using an unbiased approach. The four DS-specific clusters showed similar patterns as those of the Ctl group. The number of accessible peaks were 8,517 at D0 (DS-iPSC), 4,317 at D19 (DS-NPC), 3,579 at D35 (DS-imN) and 3,483 at D65 (DS-GABA) (**Figure 2B, 2C, Supplementary** Figure 5A-D). We identified differences in the genomic distribution of the DS-iPSC clusters when compared to that in the Ctl-iPSC clusters (**Figure 1D, 2D**). In the control group, 42.0% of the peaks were located in promoter regions, whereas in the patient group, this proportion was slightly lower at 37.3%. Additionally, a slightly higher proportion of DS-iPSC specific peaks were found in intron regions (26.5%) compared to Ctl-iPSC (24.6%). The genomic features of the DS-NPC specific cluster were similar between the Ctl group and the DS group. In the DS-imN specific cluster, a lower proportion of peaks were located in promoter regions when compared to that of the Ctl group (50.8% vs. 23.1%), while a lower proportion of peaks were from intron regions in the Ctl group (27.8%) when compared to the DS group (46.2%). In the DS-GABA and Ctl-GABA specific clusters, 34.7% and 25.9% of the peaks were located in promoter regions, respectively, while 38.7% and 43.4% of the peaks were in intron regions in the Ctl and DS groups, respectively.

We then examined the enrichment of motifs for trans-acting TFs within the differential peaks of each DS-cluster at each time-point (**Figure 2E**). The DS-iPSC cluster specific peaks showed enrichment of motifs from pluripotency related TFs, such as BRN1 and OCT4, whereas TF motifs for the SOX family of TFs were enriched in both the DS-iPSC cluster and DS-NPC cluster (**Figure 2E**). Peaks at TF motifs for LHX3 and NKX, specific for GABAergic neurons were enriched only in the DS-NPC cluster but not in the DS-GABA cluster. However, the DS- GABA specific peaks were enriched at motifs for the activator protein-1 (AP-1) family of TFs, including JUN, FRA, and FOSL. Enriched motifs for the BACH2 TF were observed only in the DS-GABA specific peaks and not in the Ctl-GABA peaks. Together, these data further support that DS-iPSC, although they initially respond to our protocol for GABAergic induction with accessible motifs for the SOX family of TFs, have a disrupted trajectory into GABAergic interneurons as shown by the loss of motifs for DLX and LHX at D65.

Next, we annotated the differential accessible peaks to nearby genes and performed KEGG pathway analysis for each cluster (**Figure 2F, Supplementary Table 4**). Similar to Ctl-iPSC, the peaks of the DS-iPSC specific cluster were enriched for pathways related to pluripotency. However, and in contrast to the control group, pathways related to GABAergic synapse were enriched already in the DS-NPC specific clusters but not in the later appearing clusters specific for DS-imN and DS-GABA (**Figure 2F**). Additionally, the pathway of long-term potentiation, a hallmark for mature GABAergic cells, was specifically enriched in the DS-imN specific cluster but not in the DS-GABA clusters. Furthermore, representative annotated genes for Ctl- iPSC lines, PAX6 and GAD2 in the Ctl-NPC and Ctl-imN clusters, and GABRG3 in the Ctl- GABA cluster, exhibit a different tendency of chromatin accessibility in DS-iPSC clusters (**Figure 2G**), also shown by qPCR analysis (**Supplementary Figure 5E**). The enrichment of TF motifs and predicted pathways relevant for GABAergic interneuron appearing already in DS-NPC, but not in DS-GABA, uncovered a pattern distinct from that in Ctl-iPSC lines, that strongly suggest a perturbed GABAergic development associated with *SCN1A* variants.

### Common chromatin dynamics in Ctl- and DS-iPSC models of GABAergic development

Identification of abnormal chromatin dynamics in GABAergic interneuronal development of DS-iPSC lines requires a thorough comparison with Ctl-iPSC lines. To extract shared dynamic chromatin features in DS-patient and Ctl-iPSC lines we first merged the differentially accessible chromatin peaks specific to each of the four time points (D0, D19, D35 and D65) within the Ctl and DS patient groups, respectively. The lists of all significant peaks in the Ctl- and DS- iPSC lines, respectively, were subsequently intersected to identify the common peak regions. In total, we identified 12,256 accessible chromatin peaks that show a shared and time-point specific pattern (**Figure 3A, Supplementary Table 5**).

**Figure 3:**
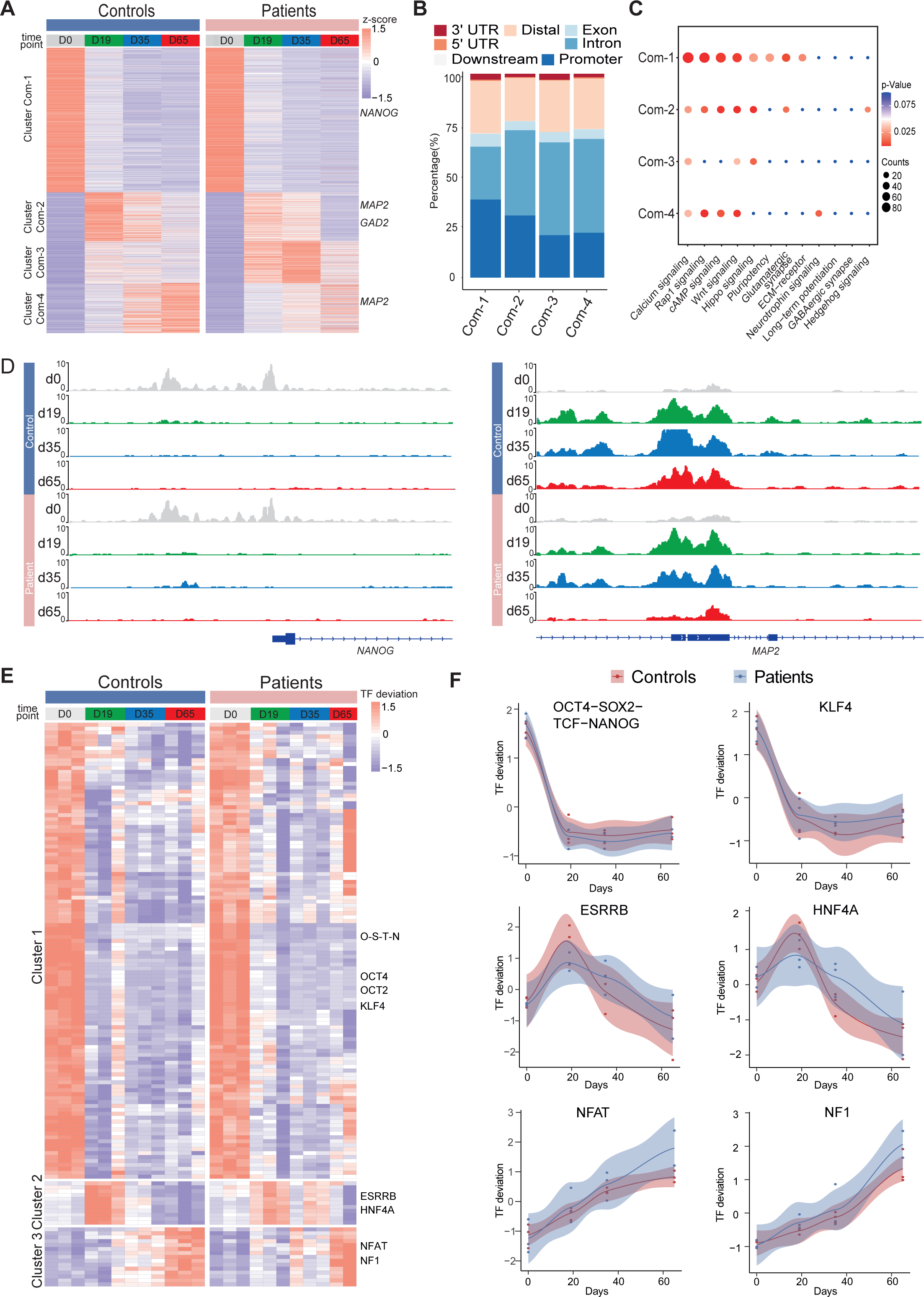
Common chromatin accessibility features of control and Dravet Syndrome patient iPSCs during GABAergic differentiation **A.** Heatmap for common chromatin accessible regions detected between control and Dravet Syndrome (DS) group during differentiation. Four clusters were obtained from time series analysis. Representative genes are labeled on the right side of heatmap. **B.** Barplot for genomic distribution of chromatin accessible regions for each cluster. **C.** Bubble plot of KEGG pathway enrichment for each cluster. p-value and enrichment were indicated. **D.** Genome browser view showing representative chromatin accessible regions at the indicated gene loci. These genes represent common changing regions between control and patients groups. **E.** Transcription factors (TFs) deviation Z scores heatmap for unique TF enriched in each cluster. Representative TFs are labeled on the right side of heatmap. **F.** Representative TFs enrichment (deviation Z score) dynamics at ATAC-Seq peaks in control and Dravet Syndrome (DS) group during differentiation.

Next, we sought to examine the genomic annotations, KEGG pathways, and TF motif enrichment for these common peaks (**Figure 3**). A K-mean clustering of read counts from the 12,252 accessible chromatin peaks revealed four common clusters (Com-1 with 6,264 peaks, Com-2 with 2,071 peaks, Com-3 with 1,793 peaks, and Com-4 with 2,124 peaks) (**Supplementary Figure 6**, **Figure 3A**). The DS-iPSC and Ctl-iPSC lines show a similar sequencing read count enrichment in Com-1 cluster and Com-2 clusters at the four time-points whereas slight differences are observed for the Com-3 and Com-4 clusters (**Figure 3A**). Accessible regions in Com-1 cluster are specifically open at D0 (iPSC) with annotations to TF genes for pluripotency such as *NANOG* and *SOX2*, among others. The Com-1 cluster remains closed at the following three time-points. The Com-2 cluster is closed at D0 but open at D19 (NPC) with accessible peaks at genes such as *MAP2*, *NEUROD1*, and *NEUROG1*, among others (**Figure 3A**).

In both the Ctl and DS-patient groups, accessible regions in the Com-3 cluster are closed D0 but slightly open at D19 (NPC) and at D35 (imN) and then more closed at D65 (GABA). However, the degree of openness for a large proportion of peaks is much higher in the DS-patient group when compared to the Ctl group. Furthermore, at D65 (GABA) the Com-3 cluster shows accessible regions at NPC-specific genes such as *MAP2* and *ATOH1* but not for regulatory elements of GABAergic neuron-specific genes, such as *ASCL1*, *LHX6*, and *DLX2*, among others. The absence of common open chromatin regions at regulatory elements for GABAergic specific genes in the Ctl and DS-patient groups supports altered dynamics of accessible chromatin at the late time-point GABAergic development in our model system.

The genomic distribution of accessible peaks with respect to promoters, exons and introns varies between the Com-clusters (**Figure 3B**). The Com-1 cluster shows a relatively large proportion of accessible chromatin regions from promoter regions (38.2%) when compared to the other clusters (Com-2: 22.0%; Com-3: 20.8%, and Com-4: 30.5%). Conversely, the Com-1 cluster shows a relatively small proportion of accessible regions in intronic regions (25.8%) when compared to Com-2, 3 and 4 clusters. However, the genomic distribution of accessible peaks was very similar in the Com-2 and Com-3 clusters.

We then conducted KEGG analysis on genes within peaks of the four Com-clusters to understand the common biological features of the Ctl- and DS-iPSC lines. We observed enrichment for the pathways of transcriptional regulation of pluripotency, that included *NANOG*, in Com-1 (**Figure 3C**) and of nitric oxide-stimulated guanylate cyclase that included *MAP2* in Com-2 (**Figure 3D**). However, in the Com-3 or Com-4 clusters, characterized by an increasing open chromatin with development, we did not find enrichment in pathways relevant for GABAergic cells (**Figure 3D**). These observations further indicate similarities in the Ctl and DS iPSC-lines in cellular processes up to D19 (NPC), but significant differences at D35 (imN) and D65 (GABA).

Furthermore, we examined the enrichment of motifs for trans-acting transcription factors (TFs) from all common peaks and clustered TFs with a k-means method. The TF enrichment patterns are similar in the Ctl and DS groups, and three TF clusters were observed. Cluster 1, containing pluripotent-specific TFs such as OCT4 and KLF, among others, shows enrichment at iPSC in both the control and DS patient groups (**Figure 3E, 3F**). Cluster 2, including TF motifs for the NPC-specific TFs ESRRB and HNF4A, is enriched in iPSC (D19) and imN (D35) (**Figure 3E, 3F**), while some TFs relevant to GABAergic cells, such as NFAT and NF1, are enriched in D65. No GABAergic neuron-specific TFs (e.g., DLX, LHX, ASCL) are observed. Taken together, our data suggest similarities between Ctl- and DS-iPSC lines in accessible chromatin at loci for TFs important for GABAergic induction and early development. However, the Ctl- and DS-iPSC groups do not show similarities in the enrichment of TFs that are critical at later stages of GABAergic maturation.

### Distinct chromatin features in DS-iPSC GABAergic development

A PCA analysis of ATAC-Seq data at D0 demonstrated that profiles of chromatin accessibility are similar in Ctl-iPSC and DS-iPSC-lines at d0 (iPSC) but different at d19 (NPC), d35 (imN), and d65 (GABA) (**Figure 4A**). These data suggest a diverging and disrupted trajectory of GABAergic development in our DS-iPSC model when compared to that in Ctl-iPSC.

**Figure 4:**
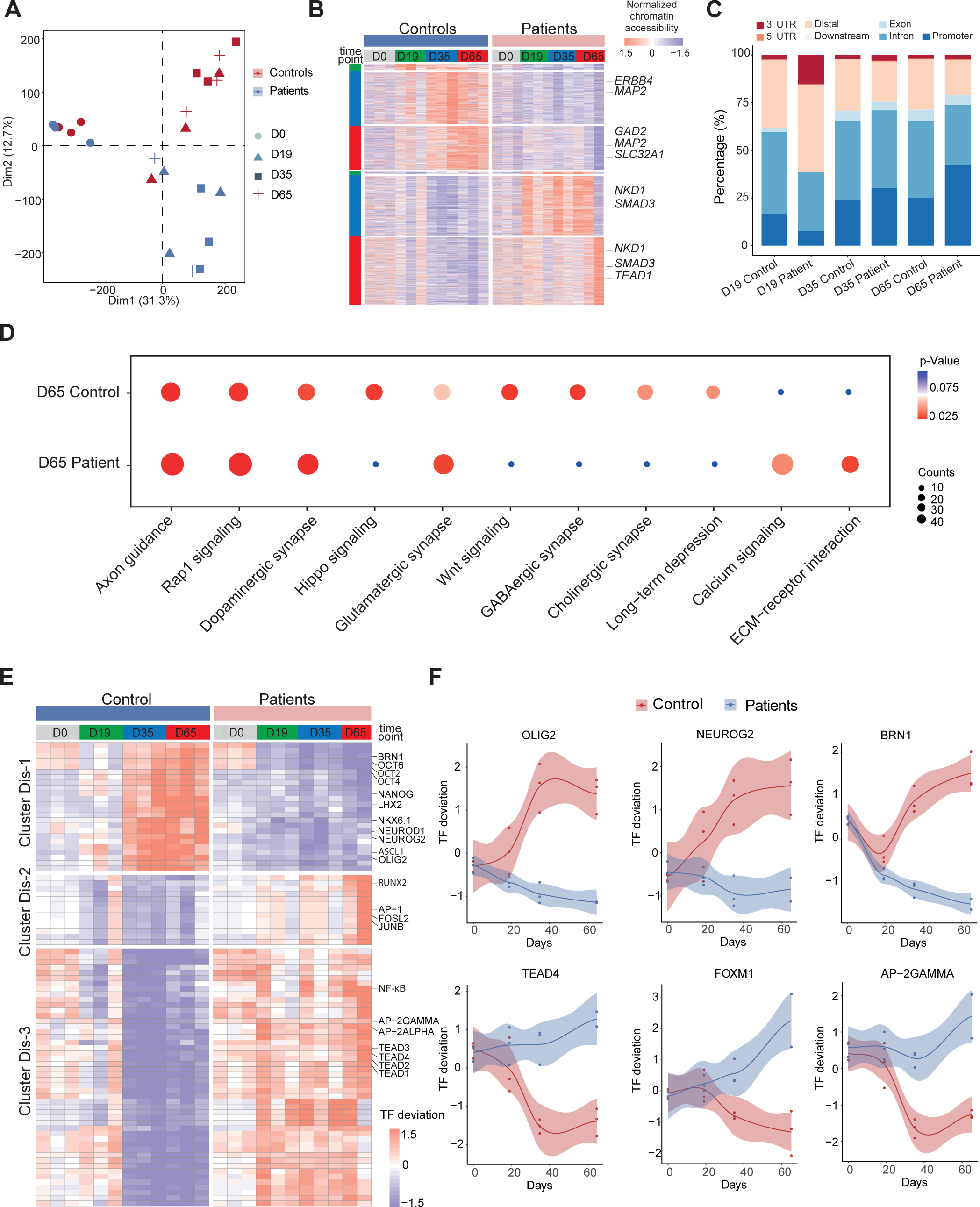
Unique chromatin accessibility features of control and Dravet Syndrome patient iPSCs during GABAergic differentiation **A.** Principal component analysis (PCA) plot of all ATAC-Seq from control and Dravet Syndrome patients during GABAergic differentiation. **B.** Heatmap showing the differential chromatin accessible peaks for comparison at each time point between the control and Dravet Syndrome (DS) patients. Representative genes are labeled on the right side of heatmap. **C.** Barplot for genomic features of unique chromatin accessible regions at each time point for the control and Dravet Syndrome (DS) patients. **D.** Bubble plot of KEGG pathway enrichment at GABAeric interneuron (Day 65) for the control and Dravet Syndrome (DS) patients. p-value and enrichment score were indicated. **E.** Heatmap showing transcription factors (TFs) enrichment at differential chromatin accessible peaks for comparison at each time point between the control and Dravet Syndrome (DS) patients. Representative TFs are labeled on the right side of heatmap. **F.** Representative TFs enrichment (deviation Z score) dynamics at ATAC-Seq peaks in control and Dravet Syndrome (DS) group during differentiation.

To further understand the molecular regulators of dysfunctional GABAergic development in DS-iPSC, we extracted the chromatin accessible peaks that were mutually exclusive in the Ctl- and DS-iPSC lines at each time point (as described in the Methods section). The peaks were categorized into a Ctl distinct peak list and a DS distinct peak list. In total, the Ctl distinct peak list contained 3,311 accessible peaks (0 peaks at D0; 42 peaks at D19; 1,490 peaks at D35; and 1,779 peaks at D65), while the DS distinct peak list contained 4,434 accessible peaks (0 peaks at D0; 13 peaks D19; 1,708 peaks D35; and 2,713 peaks D65) (**Figure 4B, Supplementary** Figure 7A, **Supplementary Table 6**). These distinct peaks were located at different genomic regions (**Figure 4C**). Furthermore, KEGG enrichment analysis of the distinct peaks at D65 revealed pathways specific to cortical neurons, such as GABAergic synapse, Glutamatergic synapse, Dopaminergic synapse, and others, in Ctl-GABA but not in DS-GABA (**Figure 4D, Supplementary** Figure 7B,).

Next, we combined the distinct peak lists of the Ctl- and DS-iPSC lines and calculated enrichment of TF motifs using a systematic approach with chromVAR (chromatin variation across regions)^39^. We then performed a non-hierarchical clustering of TFs enriched in the distinct groups and identified three clusters (Dis-1, Dis-2, Dis-3) (**Figure 4E**). Cluster Dis-1 comprised two subclusters of TFs: Subcluster 1 included the OCT family of TFs (OCT2, OCT4, OCT6) and BRN1, with enrichment restricted to D0 (iPSC) in the DS group whereas in the Ctl group the enrichment was observed at D0, D35, and D65 but not at D19. Subcluster 2 included the GABAergic specific TFs OLIGO2, ASCL1, NEUROD1, NEUROG2, NKX6.1, LHX2, and others. These TFs were not enriched at any time-point in the DS group but enriched in the control group at D35 and D65 (**Figure 4F**). In cluster Dis-2, TFs such as JUNB, AP-1, RUNX2, and others were enriched in the DS group from d0 (iPSC) to D65 (GABA). However, in the Ctl group enrichment for Dis-2 cluster was observed only at D0 (iPSC). Cluster Dis-3 is enriched in both Ctl- and DS-iPSC for TFs of the TEAD family, AP-2 and NF-kappa beta. In the Ctl group, this enrichment increased significantly in NPC and decreased (disappeared) in imN and GABAergic neurons. Interestingly, the enrichment of these TFs in cluster Dis-1 persisted in the DS group with development from NPC to GABA. The absent enrichment of GABA -specific TF motifs at D65 in the DS group brings further support for a disrupted chromatin remodeling during interneuron development.

### VPA induces changes in the chromatin landscape of iPSC derived GABAergic interneurons

Valproic acid (VPA) is a broad-spectrum anti-epileptic drug commonly used in the treatment of DS^1^. However, the response to VPA treatment varies considerable among affected individuals^40^. Prior reports suggest that the drug acts as a histone deacetylase (HDAC) inhibitor and thereby impacts chromatin remodeling^22,23^. We therefore hypothesized that chromatin changes induced by VPA in GABAergic cells may uncover processes contributing to therapeutic effects of the drug. To this end, we cultured Ctl- and DS-iPSC-derived GABAergic interneurons at D65 with and without VPA supplementation at therapeutic concentrations for six days. We then performed ATAC-Seq on three sample pairs of Ctl-GABA and on two pairs of DS-GABA (**Figure 5A, Supplementary** Figure 8A-D**, Supplementary Table 7)**. Subsequent PCA analysis of the ATAC-Seq data showed that VPA exposure induced variable changes in accessible chromatin that were seemingly iPSC-line specific (**Figure 5B, 5C, Supplementary** Figure 8E). We then focused our analysis on the sample pairs showing the most extensive VPA-induced changes in chromatin accessibility, including two Ctl-GABA pairs (Ctl1B and Ctl7C) and one DS-GABA pair (DD5A; **Figure 5C**). Genomic annotation revealed that chromatin changes induced by VPA were distributed in promoters, introns, and distal regulatory elements and at a variable extent when comparing individual Ctl and DS pairs (**Supplementary Figure 8F**). The regions showing VPA induced changes in accessible chromatin were distributed in promoters, introns, and distal regulatory elements (**Figure 5D**). The changes in chromatin accessibility after VPA treatment observed in both the Ctl- and DS- GABA may reflect unspecific effects of the drug. To validate our hypothesis on chromatin specific effects of VPA we performed enrichment analysis of differentially accessible peaks for each iPSC-GABA pair. The results show enrichment in chromatin-relevant pathways in both Ctl and DS pairs (**Figure 5E**). The analysis of open chromatin in response to VPA in Ctl- and DS-GABA thus suggest drug-induced changes that are unspecific and genome-wide. This is in line with the reported unpredictable effects of VPA on seizure frequencies in DS.

**Figure 5:**
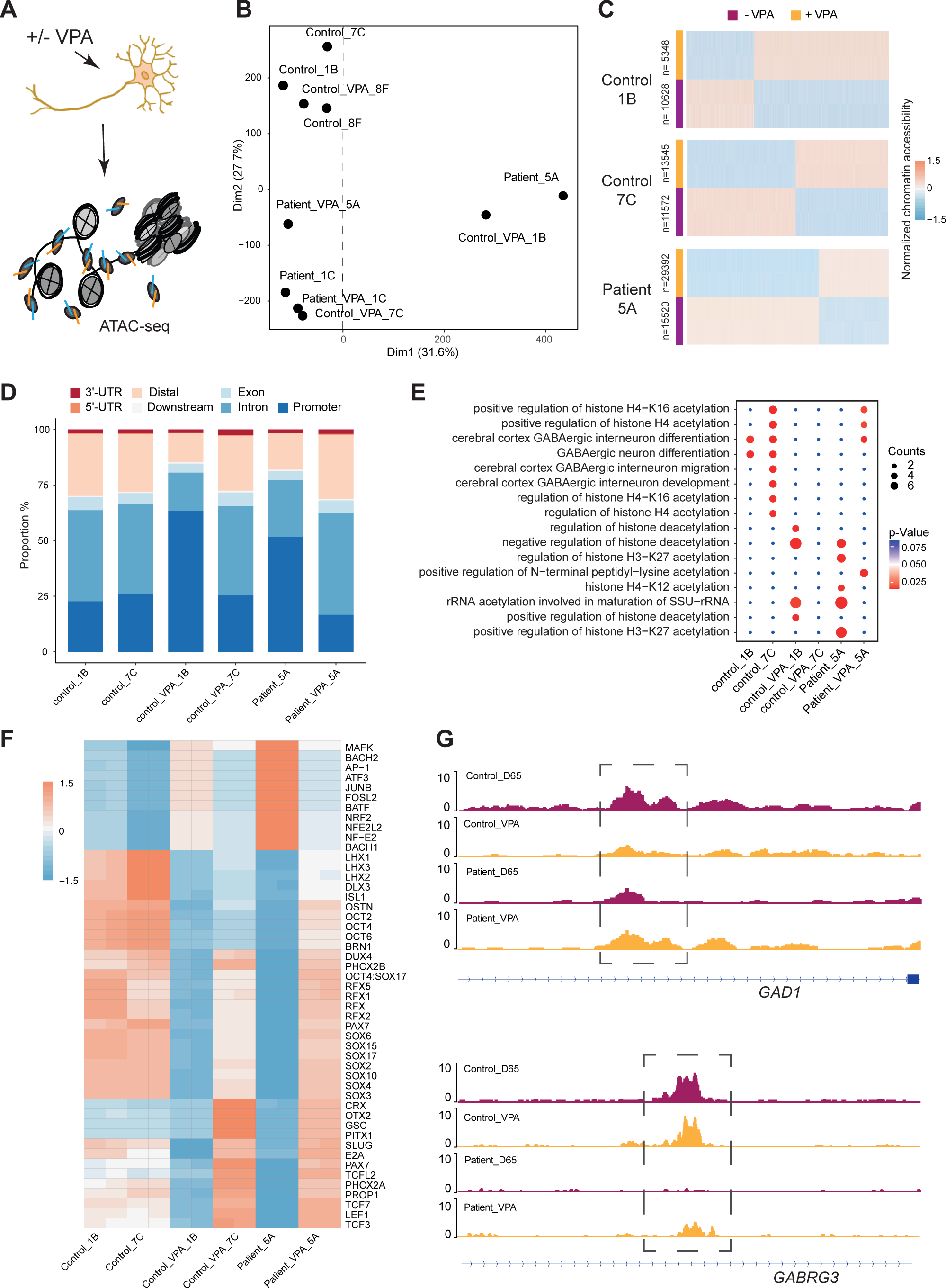
Chromatin accessibility response to Valproic acid treatment in GABAergic interneurons **A.** Schematic illustration showing measurement of chromatin accessibility response in valproic acid (VPA) treatment of GABAergic interneurons. **B.** Principal component analysis (PCA) plot of ATAC-Seq with and without VPA treatment of GABAergic interneuron in control and Dravet Syndrome patients. **C.** Heatmap of differential chromatin accessible peaks from the VPA responsible group, consisting of two control samples and one Dravet Syndrome (DS) patient sample. **D.** Barplot for genomic distribution of VPA responsible accessible regions from differential conditions. **E.** Bubble plot of KEGG pathway enrichment for each condition. p-value and enrichment were indicated. **F.** TF deviation Z score heatmap for top 50 transcription factors (TFs) enriched at responsible chromatin regions from differential conditions. **G.** Genome browser view showing representative VPA responsible chromatin accessible regions at the indicated gene loci.

Since VPA treatment has an effect on seizure frequencies in a subgroup of affected individuals, we investigated whether VPA could “restore” the chromatin accessibility profile of DS-GABA into the chromatin profile of Ctl-GABA. The pathways relevant for GABAergic cells became enriched in the chromatin regions that become accessible after VPA treatment in the DS-GABA sample pair (**Figure 5E**). A specific analysis of TF motifs becoming accessible in VPA-treated DS-GABA uncovered enriched motifs for TFs relevant for GABAergic development, such as LHX3, SOX and the RFX families of TFs (**Figure 5F**). Conversely, Ctl-GABA pairs showed an opposite enrichment pattern for these TF motifs upon VPA treatment. Furthermore, both Ctl- and DS-GABA showed a VPA-induced increase in chromatin accessibility at regulatory regions of GABAergic specific genes, such as GAD1 and GABRG3 (**Figure 5G**). Taken together, our iPSC model suggests that VPA has the potential to dramatically interfere with and change chromatin structure in GABAergic cells. These changes are likely unspecific and may affect genes important for interneuronal development.

## Discussion

The majority of cases with DS are pharmaco-resistant and there is an urgent need to improve our understanding of the underlying mechanisms for the development of targeted anti-seizure treatments ^2,40,41^. To this end, we used our iPSC-based model of GABAergic development in DS caused by heterozygous *SCN1A* variants. The model system recapitulates hallmarks of the disease and we therefore reasoned that it may improve our knowledge on DS-associated epigenetic changes as well as chromatin modifications induced by the commonly used anti-seizure drug VPA^13,29^. Specifically, we differentiated iPSCs from healthy donors (Ctl) and DS- cases towards GABAergic cortical interneurons and characterized the dynamic chromatin conformational changes using ATAC-Seq. To our knowledge, this is the first study that describes chromatin accessibility changes in DS-patient derived iPSC neurodifferentiation. We identified common and distinct chromatin accessibility profiles when comparing iPSC of DS patients and healthy donors at different time points of GABAergic development. As expected, the chromatin profiles of undifferentiated iPSC are similar in the two groups, with accessible peaks for TF genes maintaining a pluripotent state, such as *NANOG* and *POU5F1*. Differentiation towards a GABAergic fate shows that both Ctl and DS iPSC-lines acquire accessible chromatin peaks associated with genes enriched in pathways relevant for interneuronal development (e.g. signaling pathways, GABAergic synapse and long-term potentiation). However, our analyses uncovered increased changes in accessible peaks at the early phase (D19) of GABAergic development in DS-patient cells when compared to the control group. At the same time-point (D19) we also detected expression of *SCN1A*, providing a link between altered sodium flux and aberrant chromatin dynamics in DS-iPSC derived neurons. The genes associated with the distinct temporal changes of open chromatin in DS-patient cells are enriched in KEGG terms such as GABAergic synapse, Glutamatergic synapse and Long-term potentiation, suggesting a perturbed and seemingly accelerated initial GABAergic induction before D35. The enrichment in these pathways is lost with further differentiation of DS derived iPSC-lines and they fail to acquire the chromatin profile of Ctl GABAergic cells at D65. The aberrant chromatin profiles in our iPSC model comply with previous studies on iPSC and mice models of DS showing both transcriptional and electrophysiological changes in interneurons^9,13,42^. In our prior study, the transcriptional changes emerged predominantly after D19, the ATAC-Seq analysis in the current study show alterations in chromatin structure already at D19. This observation confirms that epigenetic changes detected by ATAC-Seq precede the transcriptional changes^30^. Furthermore, our study defines differential accessible chromatin regions and TF motifs associated with heterozygous and pathogenic *SCN1A* variants, bringing further insights into disturbed interneuronal differentiation in DS. In control iPSC lines, the chromatin accessibility of motifs for TFs relevant to drive neurogenesis, such as NEUROG2, OLIG2, BRN1, increase with differentiation whereas the same regions are closing in DS derived iPSC. On the other hand, motifs of more general TFs, such as TEAD4, FOXM1, AP-2GAMMA, become either accessible or un-accessible with differentiation in control iPSC but remain closed or open in differentiating DS iPSC. This is consistent with our previous observations in DS-GABA showing dysregulated expression of *FOXM1* and of genes enriched in pathways for histone modification and cell cycle pathways^13^. Moreover, similar to the *Scn1a* +/- mice model, showing an intact density of cortical and hippocampal GABAergic interneurons despite impaired functions^20^, our immunostainings revealed a comparable number and morphology of DS-patient and Ctl GABAergic neurons. These observations suggest that the proliferative capacity is relatively preserved during GABAergic interneuron development in DS.

Currently, valproic acid (VPA) is commonly used as one of the first line drugs in DS^1^ but the response on seizure frequencies is highly variable among affected cases^43,44^. The mode of action of VPA is not fully understood but it is believed to inhibit histone deacetylases, leading to changes in chromatin states and gene expression^25,45^. In our study, we aimed to study the epigenetic effect of VPA administration on iPSC derived GABAergic neurons derived from DS patients and healthy donors. The GABAergic interneurons exposed to therapeutic concentrations of VPA for six days showed extensive changes in accessible chromatin peaks in our cultures. Projection of the chromatin changes using KEGG uncovered pathways that reflect the expected mode of action of VPA, such as *histone acetylation* and *deacetylation* and *rRNA acetylation*, of importance for transcriptional activity.

However, the VPA-induced chromatin modifications in iPSC-GABA varied considerably with biological origin and the accessibility changes linked to GABAergic development were specific or shared for individual iPSC-GABA-lines. For example, in Ctl-GABA we observe an enrichment for LHX1-3 and DLX3 motifs that changes from open to closed after VPA exposure. In contrast, DS-GABA show a somewhat opposite enrichment pattern for LHX and DLX binding motifs after VPA exposure that changes from closed to open. In mice, motifs for the LHX and DLX family of TFs are enriched in the late interneuronal development that extends postnatally^19^. In addition, Ctl-GABA show a reduction in accessible SOX-binding motifs after VPA treatment whereas DS-GABA show an opposite response, acquiring a profile more similar to the untreated Ctl-GABA. The SOX-family of TFs has a global impact on both embryonic and adult neurogenesis^46^ and enrichment of SOX binding motifs are enriched in migratory interneurons^19^. These observations may suggest a mechanism by which VPA, at least in some cases, facilitates accessibility for TFs with beneficial effects on postnatal GABAergic function. Moreover, VPA treated DS-GABA uncovered enriched changes for TF motifs for AP-1 and BRN1 that made the cells acquire a pattern more similar to that in control lines. It may therefore be hypothesized that VPA induces unspecific changes in chromatin structure of iPSC-GABA that, at least in some cases, may lead to an increased accessibility for TFs linked to GABAergic development. Such a variability in response to VPA exposure in our model is consistent with the unpredictable effects of VPA on seizure-frequencies, even in cases with identical *SCN1A* variants^40^, mediated by epigenetic factors that are specific to individuals. Finally, our iPSC model of GABAergic development has limitations that should be considered. The model system applies a directed protocol during 65d that may not fully recapitulate maturation of GABAergic cells *in vivo*. Functional maturation of GABAergic interneurons occurs postnatally in humans, and models using direct differentiation from iPSC typically do not reach into cellular states comparable to postnatal cell types. More complex neuronal systems, such as 3D organoids from isogenic iPSC lines and in long-term cultures, in combination with single cell analysis, will be needed to confirm our data. Nevertheless, our findings represent an important step forward to clarify epigenetic changes in inhibitory interneurons that are associated with the progression to DS, adding data to a frame-work of knowledge for the development of targeted and personalized therapies. Further studies are now required to increase the number of independent iPSC lines with pathogenic *SCN1A* variants to validate our findings on epigenetic changes in DS and the effects of anti-seizure drugs such as VPA.

## Materials and Methods

### Cell culture, GABAergic differentiation and sampling

iPSC lines from three patients (DD1C, DD4A, DD5A)^29^ and three controls (Ctl1B, Ctl7C, Ctl8F)^47^ were cultured on human recombinant laminin LN521 (Biolamina) in Essential-8^TM^ medium (Thermo Fisher Scientific) and passaged using Trypl Express (Thermo Fisher Scientific) as described^29^. GABAergic interneuron differentiation from iPSCs was performed as previously described^13^. The protocol utilizes DUAL SMAD inhibition to induce neurogenesis towards neural stem cells for 10 days, followed by patterning with high levels of sonic hedgehog for nine days towards cortically fated neuronal progenitor cells (NPC) that mature for 46 days, i.e. a total of 65 days (Figure 1A). Neuronal cells at day 65 and onwards are viable as judged by morphological assessment by light microscopy. Differentiation was repeated at least three times per cell line.

Cell cultures were sampled at days 0 (D0), D19, D35 and D65, respectively, by harvesting cells with TryplE and collecting by centrifugation (300 x g, 3 min). Harvested cells were counted and assessed for viability using trypan blue staining and an automated EVE cell counter (NanoEntek). Samples with a viability of >90% were selected for ATAC-Seq library preparation (see below). Additionally, neuronal cell cultures at D65 were treated with valproic acid (VPA; 0,5 mM) for 6 days and subsequently harvested for analysis. Harvested cells were resuspended in 1 ml of 1 x PBS and fixed by adding an equal amount of 2% formaldehyde for 10 min at room temperature. The reaction was quenched by adding 100 ul 2,5 M glycine solution. Fixed cells were collected by centrifugation and washed twice with 1 x PBS. Cells where then resuspended in 1 x PBS, counted using an EVE automatic cell counter (NanoEntek) with Trypan Blue, and subsequently 50’000 cells/sample were processed for library preparation for ATAC sequencing.

### Tn5 transposome assembly

Tn5 transposome assembly was performed by as previously described^48^. Briefly, 2 μM annealed Tn5MErev/Tn5ME-A, 2 μM annealed Tn5MErev/Tn5ME-B, and 2 μM Tn5 transposase (Protein Science Facility at Karolinska Institutet, Stockholm) in dialysis buffer (50 mM HEPES-KOH at pH 7.2, 50 mM NaCl, 0.05 mM EDTA, 0,5 mM DTT, 0.05% Triton X-100, 5% glycerol) with 40% glycerol were mixed, incubated for 1 h at room temperature and subsequently stored at −20°C until use. The adaptor oligonucleotides (Tn5MErev: 5’-/Phos/CTGTCTCTTATACACATCT-3’, Tn5ME-A: 5′ - TCGTCGGCAGCGTCAGATGTGTATAAGAGACAG-3′ and Tn5ME-B: 5′-GTCTCGTGGGCTCGGAGATGTGTATAAGAGACAG-3′) were synthesized by Integrated DNA Technology. For oligo annealing, equimolar amounts of Tn5MErev and Tn5ME-A or Tn5ME-B and Tn5MErev, respectively, were combined in a PCR tube and annealed using the following program: 95°C for 5 min, ramp down to 25°C with −0.1°C/s increments.

### ATAC-Seq library preparation and sequencing

ATAC-Seq libraries were prepared following^48^. In brief, two technical replicates per sample containing 50,000 nuclei each were collected at 500 x g for 5 min at 4°C. Nuclei were resuspended in lysis buffer (10 mM Tris-Cl, pH 7.4; 10 mM NaCl; 3 mM MgCl2; 0.1% Igepal CA-630) and centrifuged at 500 x g for 10 min. The supernatant was removed and each nuclei pellet was resuspended in tagmentation buffer (25 μL 2 × TD buffer) with 2.5 μl Tn5 transposome and incubated at 37°C for 30 min. An equal volume of 2 x reverse crosslinking buffer (50 mM Tris-Cl, 1mM EDTA, 1% SDS, 0.2M NaCl, 5 ng/ml Proteinase K) was added to each sample and incubated at 65°C with agitation at 1200 rpm for overnight. DNA was isolated using a MinElute PCR Purification Kit (Qiagen). Sequencing libraries were prepared following described standard protocol^49^. All libraries were sequenced on an Illumina Nova-seq platform at Novogene Europe.

### RNA isolation and quantitative RT-PCR

Total RNA from iPSCs and differentiated cell populations was isolated using a miRNeasy micro kit (Qiagen). 1 μg of total RNA was reverse transcribed into cDNA using High Capacity cDNA transcription kit (ThermoFisher Scientific). Expression of marker genes was compared to expression of the two housekeeping genes GAPDH and ACTB. FastStart Universal SYBR Green Master mix (Roche) was used for qPCR using the following primers: ACTB-For: CAGGAGGAGCAATGATCTTGATCT; ACTB-Rev, TCATGAAGTGTGACGTGGACATC; GAPDH-For, GAAGGTGAAGGTCGGAGTC; GAPDH-Rev, GAAGATGGTGATGGGATTTC; SCN1A-For, TGAAGAATCCAGGCAGAAATGC; SCN1A-Rev, TCGAAATGAACGGAGAACAGA; NANOG-For, CAGCCCCGATTCTCCACCAGTCCC; NANOG-Rev, CGGAAGATTCCCAGTCGGGTTCACC; PAX6-For, AACAGACACAGCCCTCACAAACA; PAX6-Rev, CGGGAACTTGAACTGGAACTGAC; GAD2-For, AGCTGCAGCCTTAGGGATTG; GAD2-Rev, TTGCAAATGTCAGCGACAGC; GABRG3-For, TCATGGGCCTCAGAAACACC; GABRG3-Rev, CTTGCTGGCGTAGCATCTTTT. For all analyses, we used samples from three independent differentiation cultures and each sample was analyzed in triplicate. Expression was calculated as ΔCT(gene X, day Y) = CT(gene X, day Y) - CT(AVERAGE (GAPDH/ACTB, day Y)) and fold change was calculated as ΔΔCT(gene X, day Y) = ΔCT(gene X, day Y) - ΔCT(gene X, day 0) and presented as log2 [2^-(ΔΔCT)] ±SD.

### Immunofluorescence staining

Staining was performed on cells fixed with ice-cold 4% PFA and subsequently permeabilized in blocking solution (1 x PBS pH 7.4, 1% BSA, 0.1% Triton X-100). Primary antibodies against MAP2 (1:5’000; abcam, Cambridge, United Kingdom), GABA (1:1’000; Sigma, MO, United States), GAD1 (1:100, Millipore, MA, United States) and DCX (1:100, Santa Cruz, United States) were used for immunostaining and quantification. Primary antibodies were allowed to bind overnight separately or in appropriate combinations at 4℃. After washing three times in 1 x PBS, the secondary antibodies donkey anti-goat IgG AlexaFluor 633, donkey anti-rabbit IgG AlexaFluor 568 or donkey anti-mouse IgG AlexaFluor 488 (1:1’000; ThermoFisher Scientific, Waltham, MA, United States) were applied alone or in appropriate combinations for 1.5 h at room temperature in the dark. Visualization was performed on a Zeiss 510 confocal microscope (Carl Zeiss Microscopy, Jena, Germany) using Zen 2009 imaging software. Image processing was carried out using FIJI software.

### Bioinformatic and statistical analyses

#### ATAC-Seq data processing and quality analysis

After the Adapter sequence trimming, the ATAC-Seq sequencing reads were mapped to genome hg38 using bowtie2^50^. Mapped paired reads were corrected for the Tn5 cleavage position by shifting +4/-5 bp depending on the read strand. All mapped reads were extended to 50 bp centered around the Tn5 offset. PCR duplicates were removed using Picard (http://broadinstitute.github.io/picard/), and sequencing reads from chromosome M were filtered out. The Peak calling of each ATAC-Seq library was performed with MACS2^51^ with parameters -f BED, -g hs, -q 0.01, --nomodel, --shift 0. Peaks were merged into a matrix using bedtools merge ^52^. Raw reads within peaks were normalized using EdgeR’s cpm function ^53^. Log transformation was applied to these normalized peaks to calculate the Pearson correlation among duplicates. Differential ATAC peaks for clusters were selected using DESeq2^54^ with the following cutoffs: false discovery rate (FDR) < 0.05, |log2 fold change| > 1, and peak average intensity > 16. FDR values are Benjamini-Hochberg procedure corrected per default settings.

### Chromatin accessibility dynamic across differentiation

To avoid capturing the dynamic changes of accessible regions caused by variability across individuals, we initially compared the dynamic changes of chromatin accessibility cell line by cell line across differentiation. Subsequently, we extracted the common changes observed across different cell lines at each time point. Cluster analysis was conducted on both control and patient groups of ATAC-Seq data at different time points during differentiation (D0, D19, D35, and D65). The raw data were processed with background correction and normalization using Reads Per Kilobase per Million mapped reads (RPKM). Clustering analysis was performed separately for the control and patient groups of ATAC-Seq data using the Mfuzz package^55^, with minimum standard deviation and Z score parameters set at 1 and 0.5, respectively. Clusters were assigned based on the chromatin accessibility patterns of differential chromatin accessible peaks. Subsequently, clusters exhibiting significant changes were selected for further analysis.

### Annotation of unique ATAC peaks to genes and KEGG pathway analysis

Genomic annotation of each ATAC-Seq peak to its nearest gene for the differential accessible regions was done using ChIPseeker^56^. A gene promoter region was defined as 3 kb upstream and 3 kb downstream of the transcription start site. The peaks were annotated to their nearest gene within a 10 kb distance from the transcription start site (TSS). Subsequently, Clusterprofiler^57^ was used to perform Kyoto Encyclopedia of Genes and Genomes (KEGG) pathway enrichment analysis, and the results were ranked by false discovery rate (FDR). An FDR of less than 0.05 was set as significant.

### Transcription factors (TFs) motif enrichment analysis

TF motif enrichment was performed with HOMER ^58^ using the differential accessible regions as input. For TF binding prediction, chromVAR ^39^ was employed. In brief, the Homer vertebrate TF database was used as input for TF motifs in chromVAR, and then TF accessibility deviation values were calculated for each sample across the entire sample set. TF deviations with a threshold greater than 2 were retained, and TF motifs with a positive correlation with one group/cluster were selected to represent that group/cluster. TFs were ranked based on their variability within each group/cluster, and z-scores of deviations from each TF were visualized in a heatmap.

## Author contributions

Conceptualization: JS, ND, XC; Methodology: JS, XL, YD, JK, AW; Analysis and Investigation: JS, XL, YD, JK, ND, XC; Data curation and Visualization: JS, XL, YD, JK; Writing – original draft: JS, XC; Writing – Review and Editing: JS, XL, JK, YD, ND, XC; Supervision: ND, XC; Funding Acquisition: ND, XC.

## Acknowledgements

Research in the lab is funded by the Swedish Research Council (2020-01947 to N.D. and 2022- 00658 to X.C.), Hjärnfonden (FO2020-0171 and FO2022-0042 to ND), Swedish Cancer Foundation (21 1449Pj, 22 0491 JIA to X.C.), Stiftelsen Margarethahemmet (to N.D.) and Sävstaholm Society (to J.S.), Wallenberg Academy Fellow from Knut and Alice Wallenberg foundation (2023.0046 to X.C.), Uppsala University and Science for Life Laboratory. The funders played no role in study design, data acquisition and interpretation or decision to publish.

## Availability of the data

The ATAC-Seq data generated for this study have been deposited in the Sequence Read Archive (SRA) with the following number: PRJNA1013337.

---

**Supplementary Figure 1:**
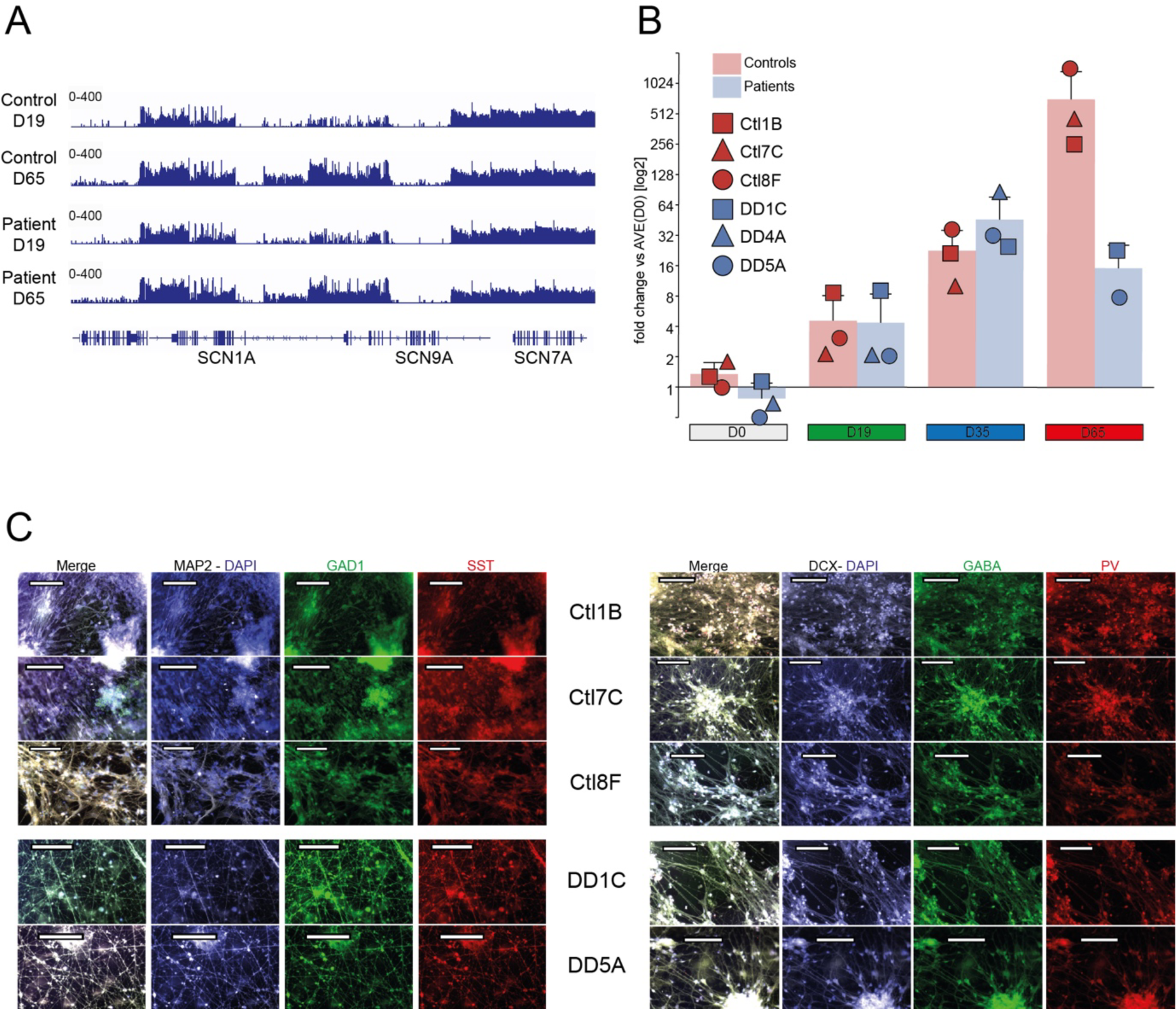
validation of GABAergic differentiation. **A.** *SCN1A* gene expression at day 19 and day 65 from RNAseq. **B.** *SCN1A* expression levels quantified by qRT-PCR following GABAergic differentiation of control and patient iPSC relative to expression of the housekeeping genes ACTB and GAPDH at D0, D19, D35 and D65. Fold change data are presented as log2 [mean ΔΔCT(day Y)] ± SD and individual data points for each cell line are indicated (Control lines in red: Ctl1B - square, Ctl7C - triangle, Ctl8F – circle; patient lines in blue: DD1C - square, DD4A - triangle, DD5A – circle; please see Methods for detail). Ctl1B, Ctl7C, Ctl8F, DD1C, DD4A, DD5A are names of cell lines. **C.** Representative pictures of immunofluorescent staining of Ctl-iPSC and DS-iPSC derived GABAergic neurons at D65 (Ctl-GABA) of GABAergic interneuron differentiation. The GABA cells form networks and stain positive for the neuronal markers MAP2 and double cortin (DCX), the GABAergic marker γ- aminobutyric acid (GABA) and the interneuronal markers somatostatin (SST), parvalbumin (PV) and glutamate decarboxylase 1 (GAD1). Size bar 100 μm.

**Supplementary Figure 2:**
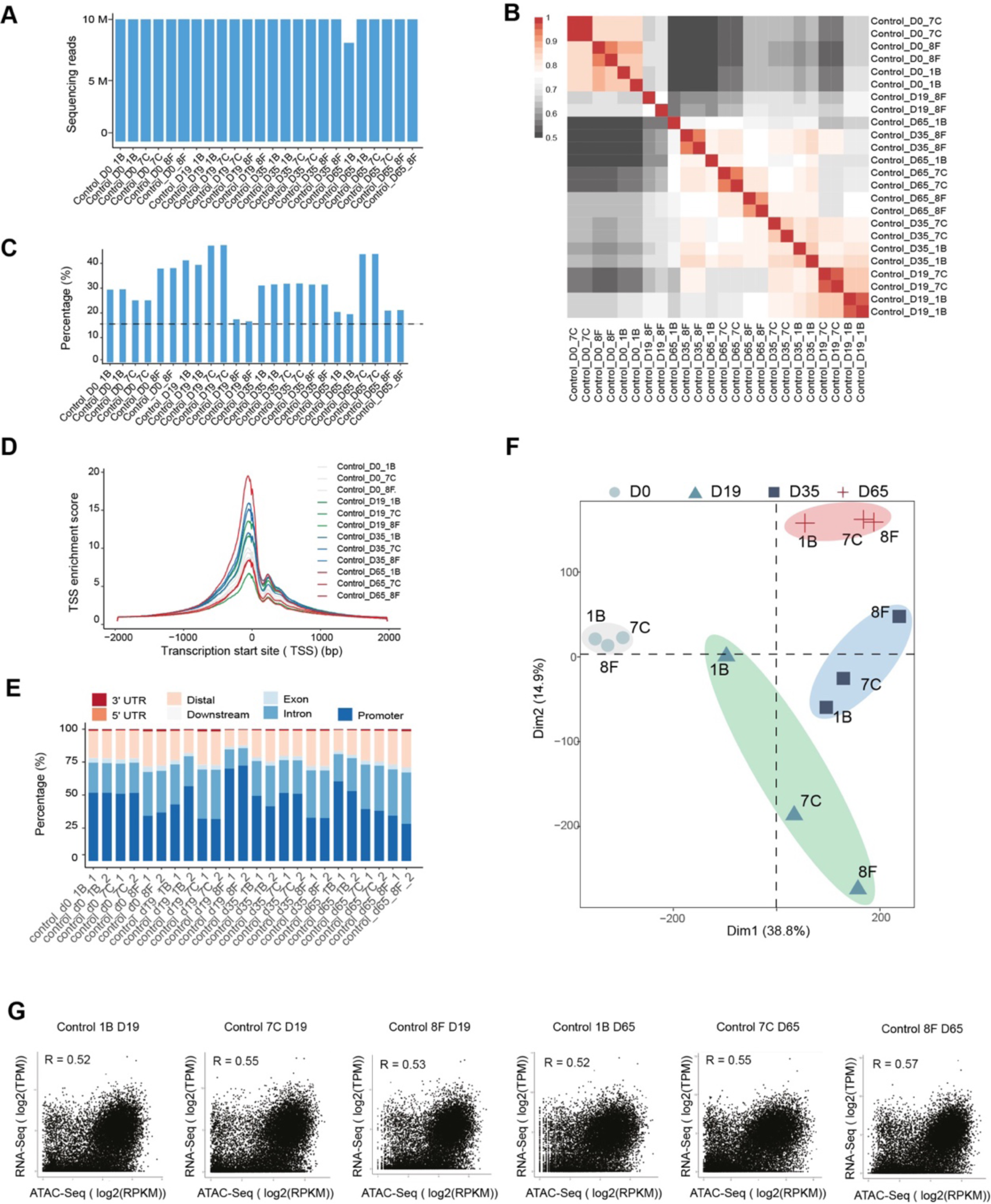
Quality control of ATAC-Seq samples during GABAergic interneuron differentiation of Ctl-iPSC. **A.** Sequencing depth for each sample. **B.** Pearson correlation matrix from technical duplicates of ATAC-Seq at different time points. **C.** Quantification of fraction of reads in peaks (FRiP) of ATAC-Seq in each condition. The dotted line indicates the cutoff used. **D.** Transcription start site (TSS) enrichment score of ATAC-Seq in each sample. **E.** Genomic distribution of chromatin accessible peaks for each sample. **F.** Principal component analysis of ATAC-Seq from each sample. **G.** Pearson correlation of gene expression from RNA-Seq and chromatin accessibility from ATAC-Seq.

**Supplementary Figure 3:**
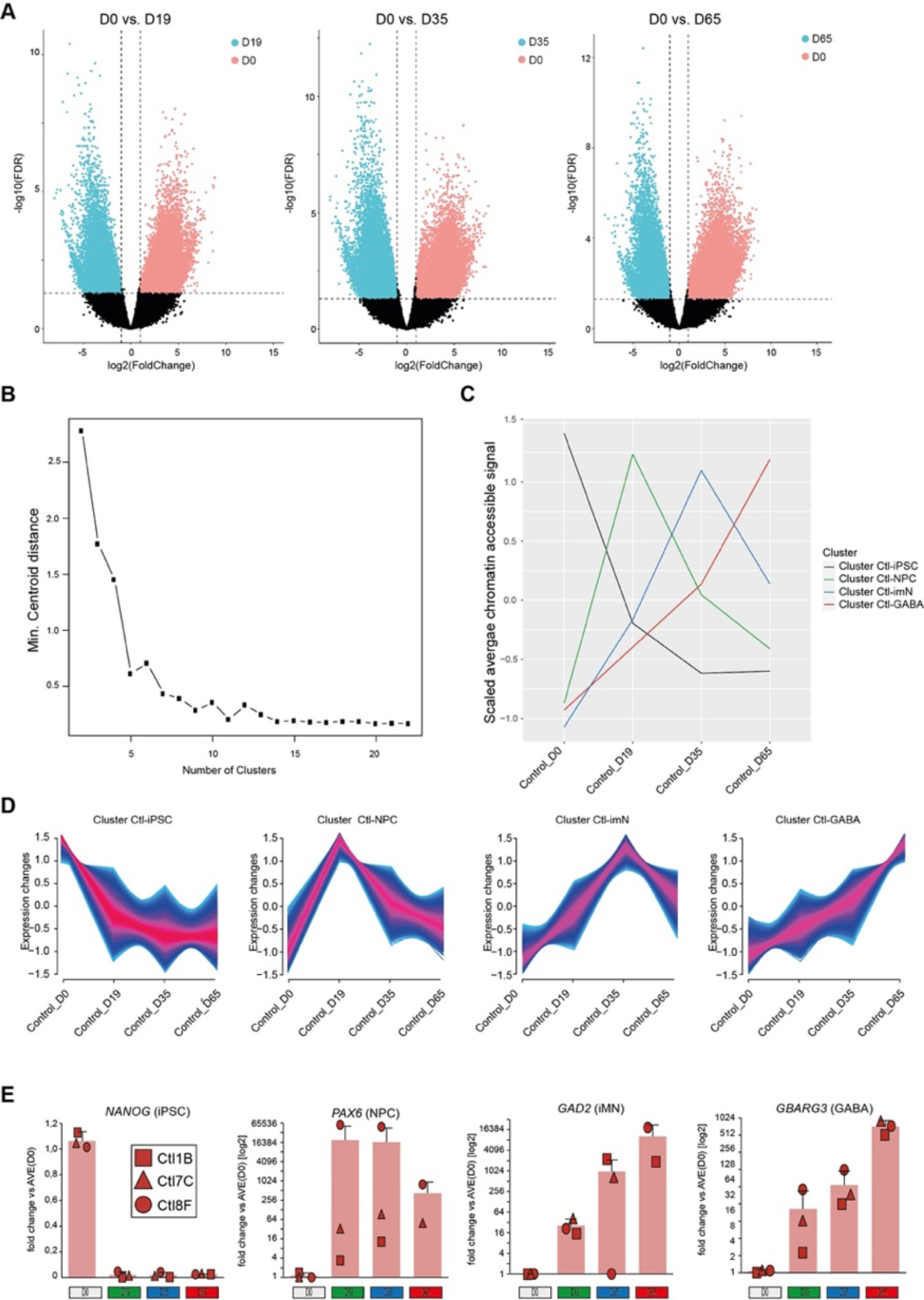
Chromatin accessibility cluster identification during GABAergic interneuron differentiation of Ctl-iPSC. **A.** Volcano plots show differential accessible chromatin peaks between D0 and different time points. D0 (n = 6046) vs. D19 (n = 3866); D0 (n = 8880) vs. D35 (n = 7865); D0 (n = 7870) vs. D65 (n = 7776). The dotted lines indicate the cutoff used to identify differential ATAC-Seq peaks. **B.** Variance with different numbers of clusters derived from chromatin accessibility during GABAergic interneuron differentiation **C.** The scaled average chromatin accessibility during GABAergic interneuron differentiation from each cluster. **D.** The tendency of chromatin accessibility during GABAergic interneuron differentiation from each cluster. **E.** Expression of the four cluster markers *NANOG* (iPSC; D0), *PAX6* (PNC; D19), *GAD2* (imN; D35) and *BABRG3* (GABA; D65) quantified by qRT-PCR following GABAergic differentiation of control iPSC relative to expression of the housekeeping genes ACTB and GAPDH. Fold change data are presented as log2 [mean ΔΔCT(day Y)] ± SD and individual data points for each cell line are indicated (Control lines in red: Ctl1B - square, Ctl7C - triangle, Ctl8F – circle please see Methods for detail).

**Supplementary Figure 4:**
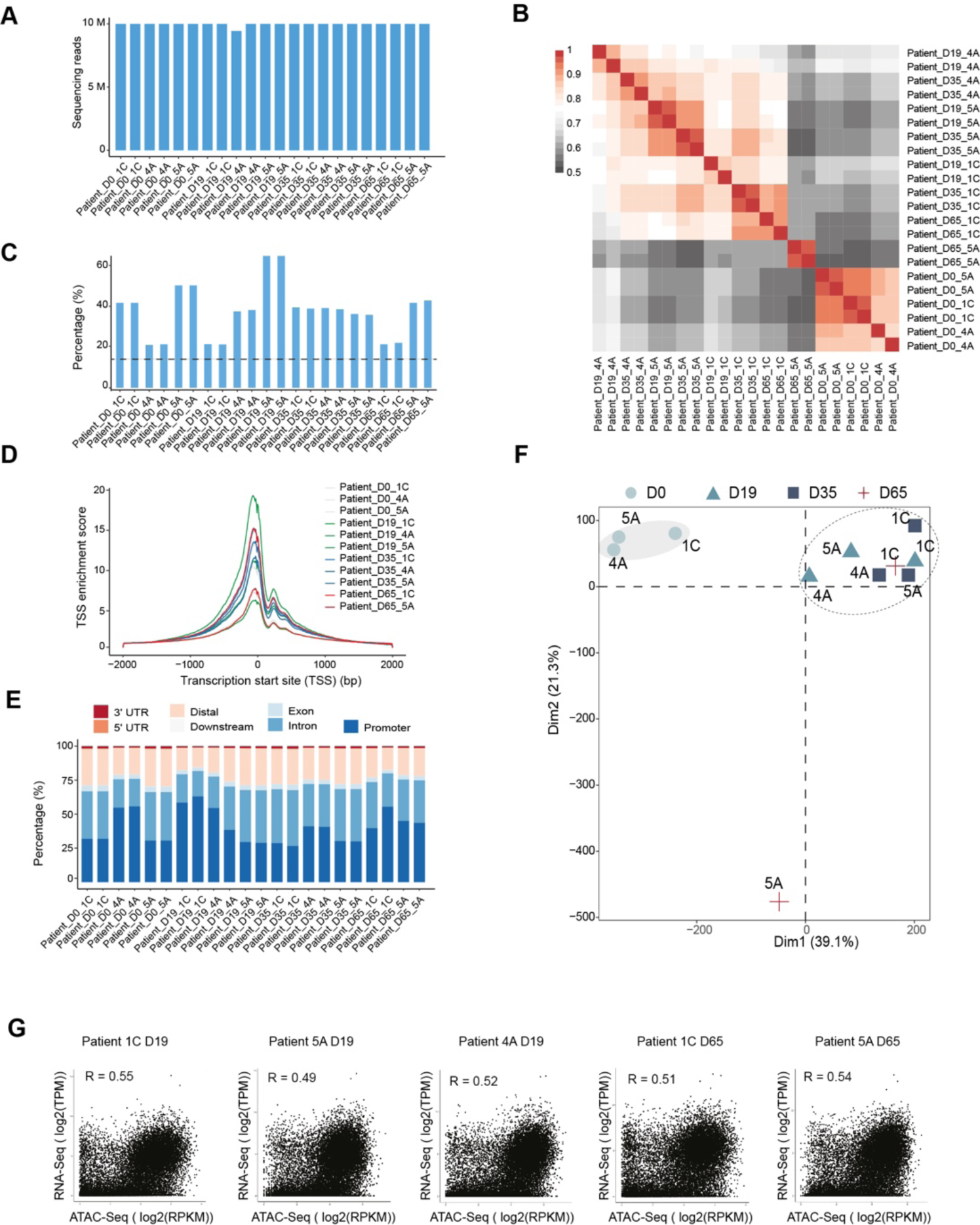
Quality control of ATAC-Seq samples during GABAergic interneuron differentiation in Dravet Syndrome patient iPSCs. **A.** Sequencing depth comparison from each sample. **B.** Pearson correlation matrix from technical duplicates of ATAC-Seq from all samples. **C.** Quantification of fraction of reads in peaks (FRiP) of ATAC-Seq in each condition. The dotted line indicates the cutoff was used. **D.** Transcription start site (TSS) enrichment score of ATAC-Seq in each sample. **E.** Genomic distribution of chromatin accessible regions for each sample. **F.** Principal component analysis of ATAC-Seq from each sample. **G.** Pearson correlation of gene expression from RNA-Seq and chromatin accessibility from ATAC-Seq.

**Supplementary Figure 5:**
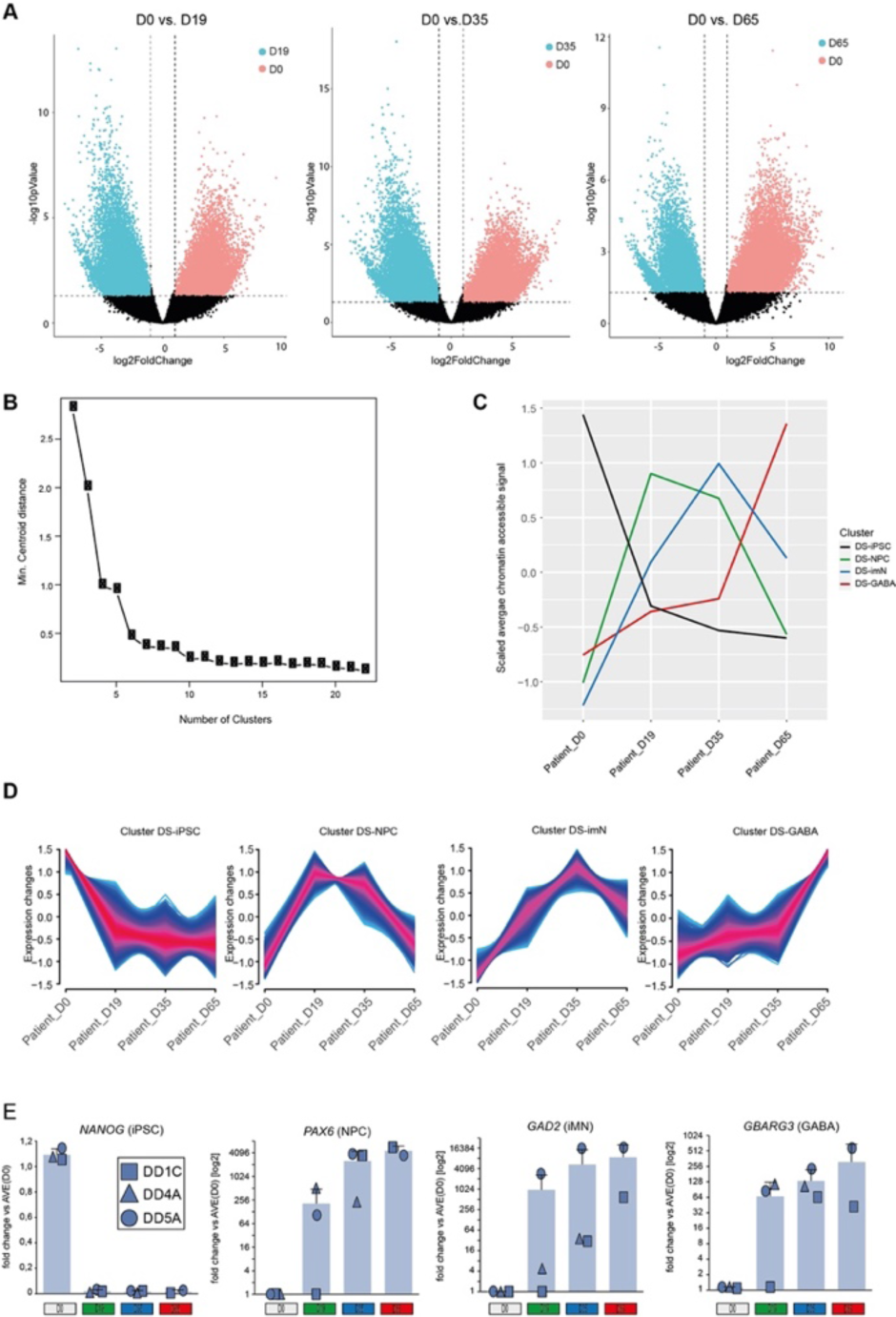
Chromatin accessibility cluster identification during GABAergic interneuron differentiation in Dravet Syndrome patient iPSCs. **A.** Volcano plots show differential accessible chromatin peaks between D0 and different time points. D0 (n = 7798) vs. D19 (n = 5183); D0 (n = 10154) vs. D35 (n = 7239); D0 (n = 6909) vs. D65 (n = 5667). The dotted lines indicate the cutoff used to identify differential ATAC-Seq peaks. **B.** Variance with different numbers of clusters derived from chromatin accessibility during GABAergic interneuron differentiation in Dravet Syndrome patients. **C.** The scaled average chromatin accessibility during GABAergic interneuron differentiation from each cluster in Dravet Syndrome patients. **D.** The tendency of chromatin accessibility during GABAergic interneuron differentiation from each cluster in Dravet Syndrome patients. **E.** Expression of the four cluster markers *NANOG* (iPSC; D0), *PAX6* (PNC; D19), *GAD2* (imN; D35) and *BABRG3* (GABA; D65) quantified by qRT-PCR following GABAergic differentiation of control iPSC relative to expression of the housekeeping genes ACTB and GAPDH. Fold change data are presented as log2 [mean ΔΔCT(day Y)] ± SD and individual data points for each cell line are indicated (DS lines in blue: DD1C - square, DD4A - triangle, DD5A – circle please; see Methods for detail).

**Supplementary Figure 6:**
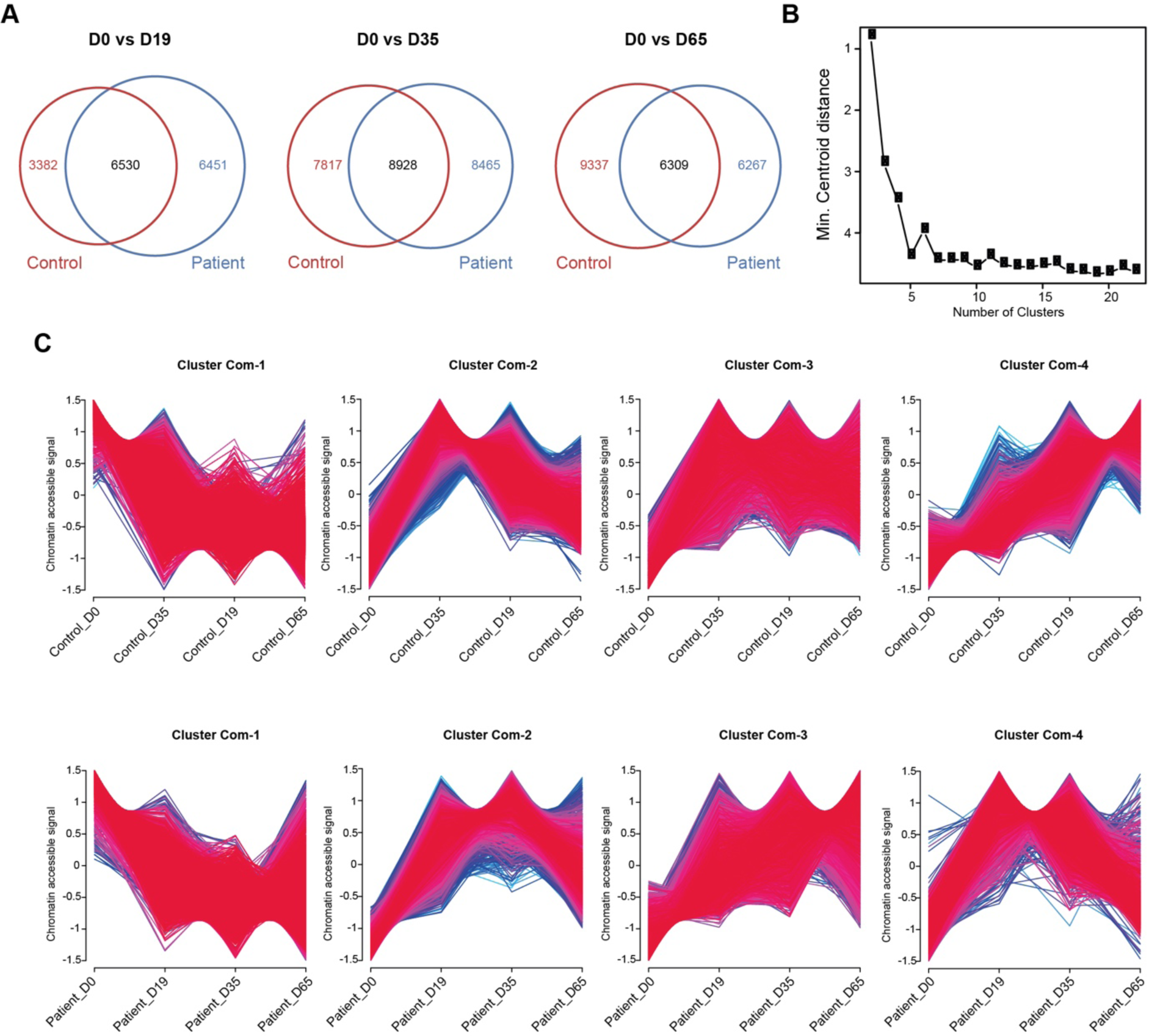
Identification of common chromatin accessibility between control and Dravet Syndrome patient iPSCs during GABAergic interneuron differentiation. **A.** Venn diagrams showing the shared and distinct accessible chromatin peaks between control and patient samples. **B.** Variance with different numbers of clusters derived from common accessible chromatin regions between control and Dravet Syndrome patients. **C.** The tendency of chromatin accessibility from common accessible chromatin regions between control and Dravet Syndrome patients during GABAergic interneuron differentiation.

**Supplementary Figure 7:**
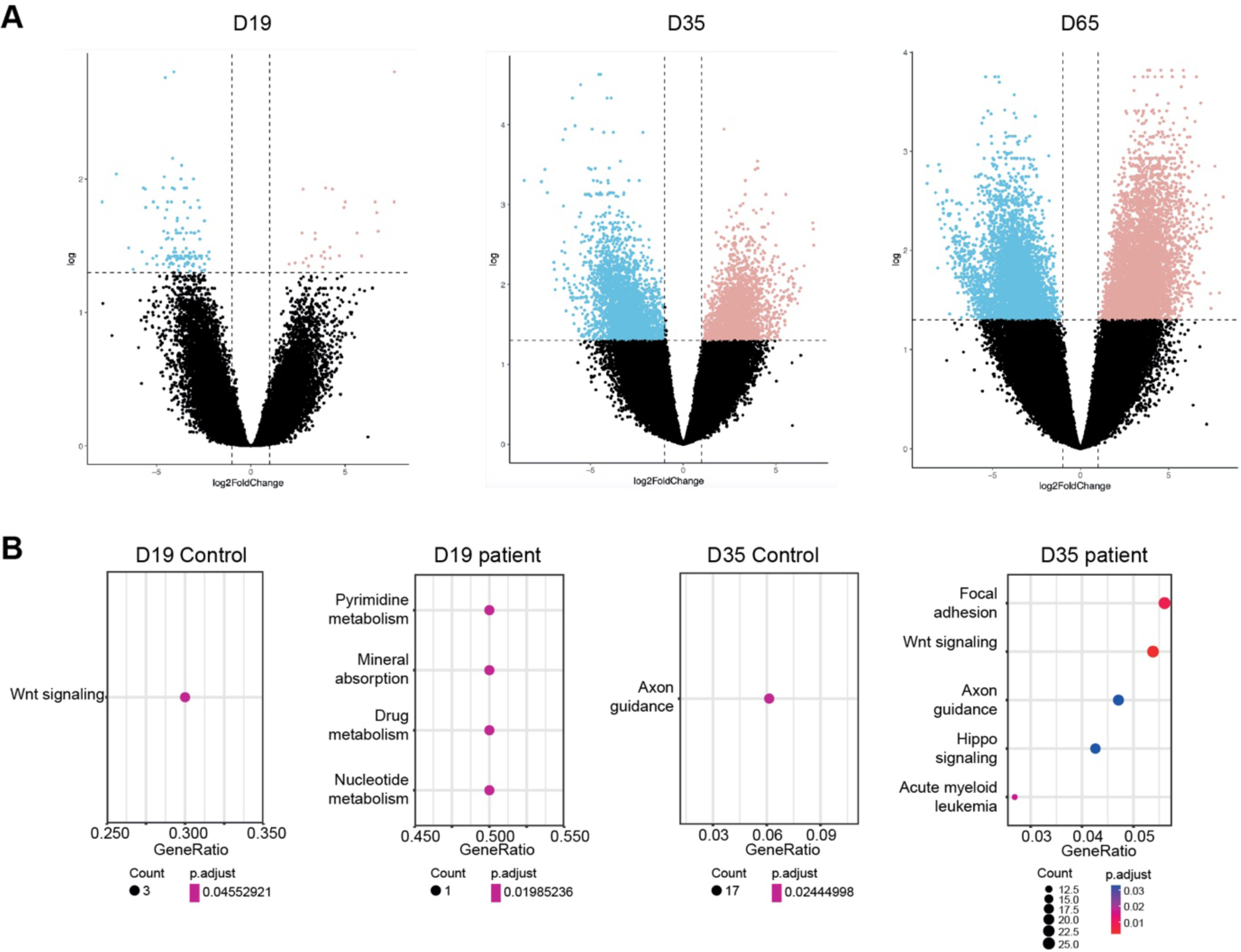
Identification of unique accessible chromatin region between control and Dravet Syndrome patient iPSCs during GABAergic interneuron differentiation. **A.** Volcano plot of differential accessible chromatin peaks between control and Dravet Syndrome (DS) patients at different stage of differentiation (NPC (Day 19), imN (Day 35) and GABAergic interneuron (Day 65)). Dotted lines indicate cutoff from fold change and false discovery rate. **B.** KEGG signal pathway enrichments from unique accessible chromatin region in control and Dravet Syndrome patients at NPC (Day 19), imN (Day 35).

**Supplementary Figure 8:**
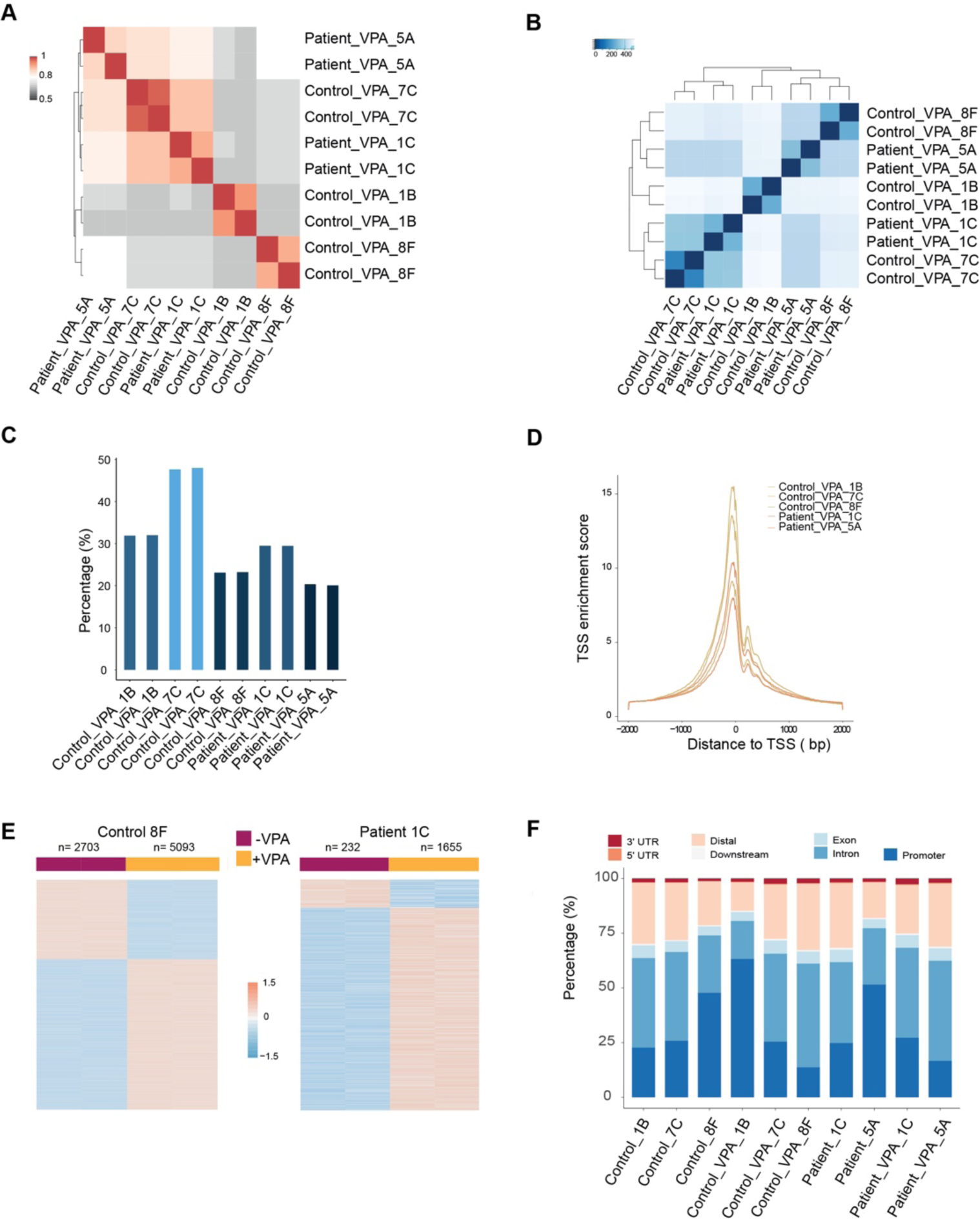
Valproic acid treatment reshapes chromatin accessibility of GABAergic interneurons. **A.** Pearson correlation matrix from technical duplicates of ATAC-Seq of VPA treated control and Dravet Syndrome patient group. **B.** Distance matrix of ATAC-Seq from VPA treated control and Dravet Syndrome patient group. **C.** Quantification of fraction of reads in peaks (FRiP) of ATAC-Seq in each condition. **D.** Transcription start site (TSS) enrichment score of ATAC-Seq in each sample. **E.** Heatmap of differential chromatin accessible regions from VPA non-responsible group, consisting of one control sample and one Dravet Syndrome (DS) patient sample. **F.** Genomic distribution of chromatin accessible regions for each individual sample.

Supplementary Table 1: Identified differential ATAC-Seq peaks during GABAergic interneuron differentiation of Ctl-iPSC.

**Supplementary Table 2: KEGG enrichment for each cluster during GABAergic interneuron differentiation of Ctl-iPSC.**

**Supplementary Table 3: Identified differential ATAC-Seq peaks during during GABAergic interneuron differentiation in Dravet Syndrome patient iPSCs.**

**Supplementary Table 4: KEGG enrichment for each cluster during GABAergic interneuron differentiation in Dravet Syndrome patient iPSCs.**

**Supplementary Table 5: Identified common ATAC-Seq peaks during GABAergic interneuron differentiation between control and Dravet Syndrome patient iPSCs.**

**Supplementary Table 6: Identified distinct ATAC-Seq peaks during GABAergic interneuron differentiation from control and Dravet Syndrome patient iPSCs.**

**Supplementary Table 7: Identified differential ATAC-Seq peaks between with and without VPA treatment.**

